# Modified Chlorophyll Pigment at Chl_D1_ Tunes Photosystem II Beyond the Red-Light Limit

**DOI:** 10.1101/2024.07.13.603357

**Authors:** Friederike Allgöwer, Abhishek Sirohiwal, Ana P. Gamiz-Hernandez, Maximilian C. Pöverlein, Andrea Fantuzzi, A. William Rutherford, Ville R. I. Kaila

## Abstract

Photosystem II (PSII) is powered by the light-capturing properties of chlorophyll *a* pigments that define the spectral range of oxygenic photosynthesis. Some photosynthetic cyanobacteria can acclimate to growth in longer wavelength light by replacing five chlorophylls for long wavelength pigments in specific locations, including one in the reaction center (RC). However, the exact location and the nature of this long wavelength pigment still remain uncertain. Here we have addressed the color-tuning mechanism of the farred light PSII (FRL-PSII) by excited state calculations at both the *ab initio* correlated (ADC2) and linear-response time-dependent density functional theory (LR-TDDFT) levels in combination with large-scale hybrid quantum/classical (QM/MM) simulations and atomistic molecular dynamics. We show that substitution of a single chlorophyll pigment (Chl_D1_) at the RC by chlorophyll *d* leads to a spectral shift beyond the far-red light limit, as a result of the protein electrostatic, polarization and electronic coupling effects that reproduce key structural and spectroscopic observations. Pigment substitution at the Chl_D1_ site further results in a low site energy within the RC that could function as a sink for the excitation energy and initiate the primary charge separation reaction, driving the water oxidation. Our findings provide a basis for understanding color-tuning mechanisms and bioenergetic principles of oxygenic photosynthesis at the far-red light limit.

## INTRODUCTION

Photosystem II (PSII) is a light-driven water oxidizing and plastoquinone reducing enzyme that transduces sunlight into chemical energy.^1^ The charge separation generates a proton motive force (*pmf*) across the photosynthetic membranes that drives the synthesis of ATP^2, 3^ and biomass, whilst the release of oxygen powers aerobic life.^4^ The photochemistry of PSII occurs in its reaction center (RC), comprising four chlorophyll and two pheophytin pigments, symmetrically embedded in the D1 and D2 proteins (Figure 1A). These RC pigments comprise a central P_D1_/P_D2_ chlorophyll pair together with two adjacent chlorophylls (Chl_D1_, Chl_D2_) and two pheophytin pigments (Pheo_D1_, Pheo_D2_) (Figure 1B).^5, 6^ This pigment array traps the excitation energy transferred from the light-harvesting antenna and functions as the site for the light-driven charge separation,^6^ driving water oxidation at the manganese-oxo-calcium cluster (Mn_4_O_5_Ca).

**Figure 1.**
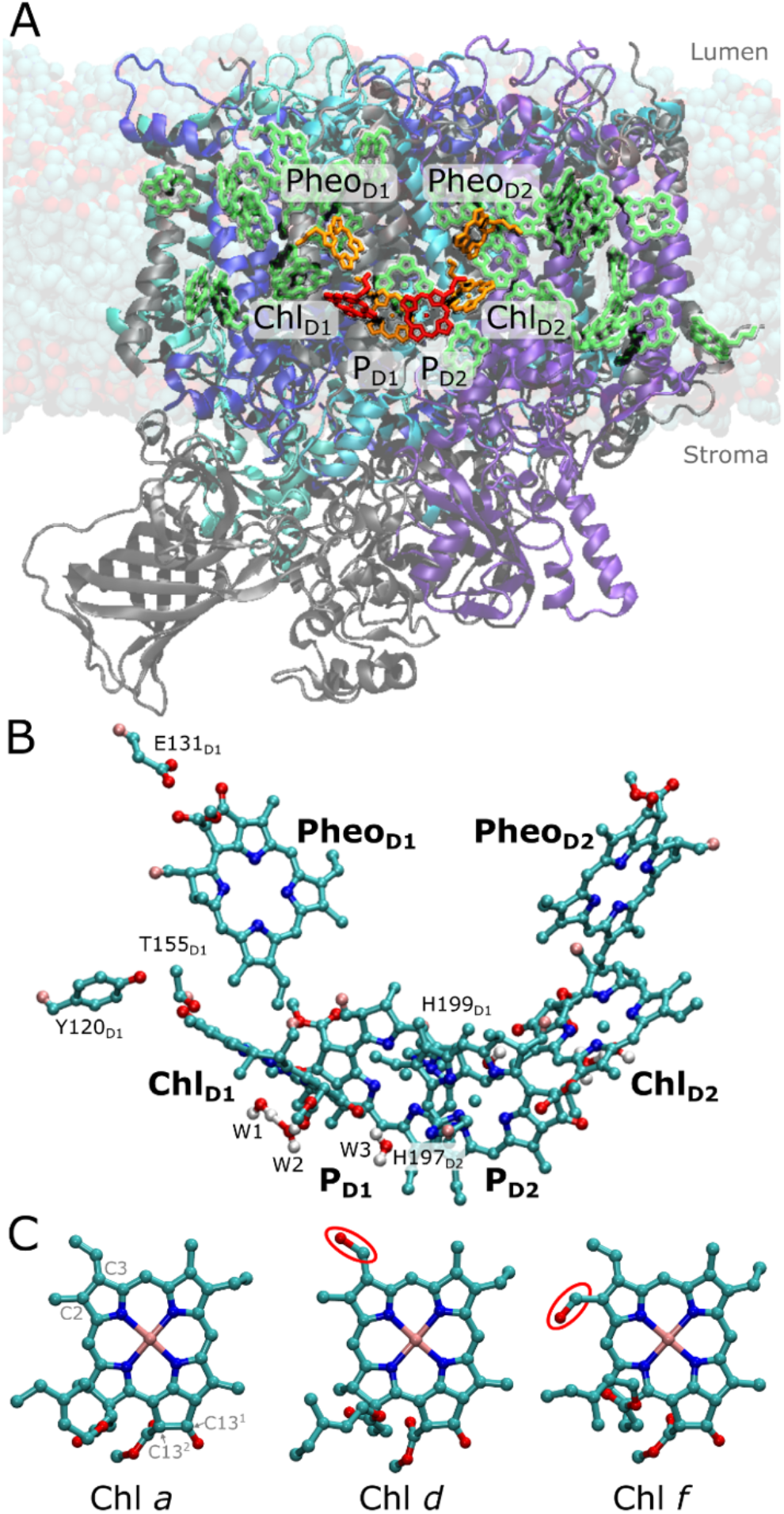
Structure of FRL-PSII. (A) Atomistic model of FRL-PSII embedded in a lipid bilayer. The modified protein segments relative to the WL-PSII are shown in blue colors, while unmodified protein is shown in gray. The chlorin-based pigments within the RC are highlighted in orange. Four models of the FRL-PSII were created by modeling Chl *d* or Chl *f* at either the Chl_D1_ or P_D2_ site (in red). (B) Closeup of the RC, involving six pigments, hydrogen-bonding residues and nearby water molecules. P_D1_ is axially ligated by D1-His198 in the WL-PSII and D1-His199 in the FRL-PSII, while P_D2_ is axially ligated by D2-His197. Chl_D1_ is axially coordinated by a water molecule (W1), which further interacts through hydrogen-bonding interaction with another water (W2). The figure also shows the W3 water molecule interacting with the C13_1_ keto group of Chl_D1_ and the axial histidine ligand of P_D2_. Chl_D2_ is similarly surrounded by three water molecules. Hydrogen atoms are not shown for clarity. (C) Comparison of the structure of different chlorophyll pigments: Chl *a* has vinyl group at the C3 position, while Chl *d* features a formyl group at the C3 position and Chl *f* a formyl at the C2 position. The differences in structure are highlighted with a red circle.

Chlorophyll *a* (Chl *a*) pigments, abundantly present in the photosynthetic proteins, enable efficient lightharvesting of PSII with a maximum absorption around 670 nm (1.85 eV). It has been suggested that the specific Chl *a* involved in PSII photochemistry at 685 nm (1.81 eV) define the “red-limit” of oxygenic photosynthesis, where photons beyond 700 nm (energy < 1.77 eV) do not efficiently trigger the charge separation responsible for water oxidation and release of O_2_. However, the cyanobacterium *Chroococcidiopsis thermalis* acclimatizes to far-red light, by replacing its conventional PSII (white-light PSII; WL-PSII) with a far-red isoform (FRL-PSII), which incorporates one Chl *d* and four Chl *f* pigments instead of Chl *a* into specific locations, while retaining all the other Chl *a* pigments in the antenna and reaction center (FRL-PSII and FRL-PSI) (Figure 1A).^7, 8^ This site-specific ‘doping’ of the pigments in the FRL-PSII, results in a significant redshift of the absorption spectrum at room temperature with a peak absorption maximum around 715-725 nm (1.71-1.74 eV), partially resolved at cryo-temperatures.^7^

Locating the position and function of these far-red light pigments in PSII is crucial for understanding the mechanistic principles of far-red light driven oxygenic photosynthesis. Nürnberg *et al*.^7^ proposed that the farred chlorophylls were not just acting as an extension to the antenna, but crucially, one of them was serving as the primary donor within the RC. This proposal was based on the efficient far-red light-driven charge separation in both room and low temperature, as well as hyper-luminescence compared to conventional cyanobacteria PSII. The finding that the Chl *d* and *f* pigments not only worked efficiently to absorb lower energy photons, but also acted as the primary electron donor in primary charge separation set both thermodynamic and kinetic boundaries on the chemically challenging photochemistry.^7^

Structural factors can be inferred by comparing the chlorophyll binding sites in WL-PSII and FRL-PSII, in particular by the identification of amino acid changes from WL to FRL that introduce new hydrogen-bonding partners for the modified formyl group, leading to a stabilization of the Chl *d* or *f* in the binding sites. Despite previous modelling efforts, spectroscopy, and structural studies, the exact location of all the far-red light absorbing pigments remain uncertain.^7-13^ For the primary donor, Nürnberg *et al*.^7^ proposed that the Chl_D1_ site was occupied by either a Chl *f* or the Chl *d* pigment, while the remaining four Chl *f* pigments were located in the CP43 and CP47 antennae (cf. Ref. 7 for spectroscopic assignment). However, based on the assignment of low-temperature circular dichroism data Judd *et al*.^14^ discussed the possibility that the P_D2_ site could be occupied by Chl *f*, while accepting that the Chl_D1_ location was also a likely candidate. Ultrafast spectroscopic experiments^10^ were interpreted in a model consistent with a far-red pigment at Chl_D1_, the primary donor, but at that time P_D2_ was not considered as an option. Recent cryo-EM data (PDB ID: 7SA3)^12^ suggested a Chl *d* pigment at the Chl_D1_ site and four Chl *f* pigments in the integral CP43/CP47 antenna complexes, but these assignments were made at least in part based on the presence of hydrogen-bonding residues close to the putative formal group substitutions, rather than on direct detection of the formyl oxygens, which are structurally difficult to detect_11, 15_ as the pigments differ only by the chemical substituents on C2 and C3 centers (Figure 1C).

The changes in the pigments are accompanied by subtle alterations in the pigment binding pockets._7, 8, 12_ In the FRL-PSII, the D1 core protein has D1-Tyr120 and D1-Thr155 near Chl_D1_, rather than D1-Phe119 and D1Ala154 in WL-PSII (Figure S1, S2).^7^ The positioning of D1-Tyr120 and D1-Thr155 are thought to facilitate hydrogen-bonding with the formyl group at C3 and C2 of Chl *d* and Chl *f*, respectively. In addition, a substitution was observed near Pheo_D1_, which remains as a Pheo *a* in FRL-PSII, where the hydrogen-bonding D1Gln130 in the WL-PSII is replaced by D1-Glu131 in the FRL-PSII. This substitution near Pheo_D1_ is observed as a response to high-light conditions in Chl *a*-containing PSII (encoded by psbA3 gene in cyanobacteria *T. elongatus* and in chloroplast PSII) tuning its reduction potential to higher values.^16, 17^ However, the mechanisms by which these substitutions could facilitate pigment-binding and spectral tuning remain unclear.

To address the molecular basis of the spectral tuning mechanism to longer wavelengths and its bioenergetic consequences, we perform here large-scale multiscale quantum/classical (QM/MM) simulations and excited state calculations at the correlated *ab initio* quantum chemical level, using second-order algebraic diagrammatic construction (ADC(2)) theory,^18^ as well as linear-response time-dependent density functional theory (LR-TDDFT). To probe the functional dynamics of the substitutions, we also performed atomistic molecular dynamics simulation (MD) models of both the FRLPSII and WL-PSII (Figure S3, Table S1) together with electrostatic calculations to probe redox tuning effects. Our atomistic models allow us to systematically probe the effect of Chl *d* and Chl *f* substitutions at distinct positions in the RC, and to derive spectral tuning principles. Our combined findings provide a basis for understanding how the RC traps excitation energy, and triggers charge-separation using the decreased energy available in far-red light.

## RESULTS

### Probing the effect of long wavelength chlorophylls in the RC

To determine the location and nature of the long wavelength chlorophyll in the FRL-PSII RC, we studied the group of four chlorophylls pigments that are involved in charge separation and the first stabilization step, P_D1_, P_D2_, Chl_D1_, and Pheo_D1_, using QM/MM calculations (Figure 2D). First, we calculated the optical spectra when either of the far-red chlorophylls were introduced into the two candidate locations within the RC, *i.e*., Chl_D1_, and P_D2_. We note that while the focus here is on the far-red pigment within the RC, the experimental spectra reflect the properties of all chlorophyll, including the far-red substitutions also in the antenna protein. However, it has been established that the RC chlorophyll absorbs at around 720 nm, as indicated in Figure 2.^7^ We then decomposed the tuning effects into site energies and excitonic coupling effects with their neighboring pigments (see *Methods*). To this end, we calculated the vertical excitation energies (VEEs) of the RC pigment cluster using QM/MM models of the FRLPSII and WL-PSII, allowing us to account for the photoexcitation within the pigment cluster at a correlated quantum mechanical level, including the polarizing effect of the complex PSII surroundings (Figure S3).

**Figure 2.**
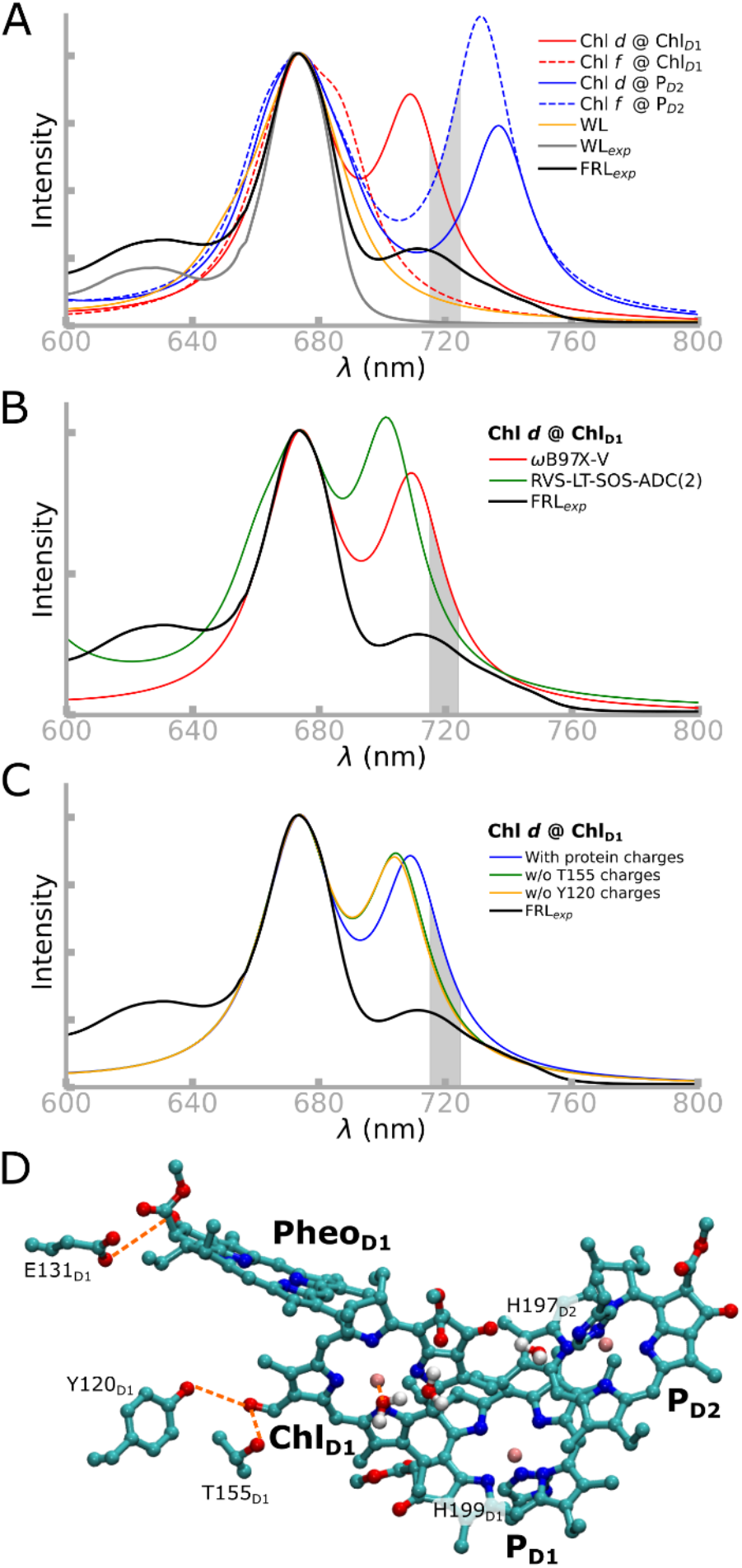
Absorption spectrum of the FRL-PSII reaction center. (A) Computed absorption spectra based on the tetramer assembly computed at the QM(TDDFT)/MM level (ωB97X-V/def2-TZVP, see *Methods*) and comparison with the experimental spectra of the FRL-PSII and WL-PSII. The VEEs and the corresponding oscillator strengths were used to compute the spectrum, which was aligned with respect to the 674 nm peak to help the comparison (see Supporting Information, *Extended Methods*). The FRL region (715-725 nm) is highlighted. Note that the experimental spectra contain the effect of substitutions also beyond the RC (see ref. 7). The corresponding ADC(2) spectra can be found in Figure S4A. (B) Closeup of the Q-band in the red and far-red region of the spectrum computed at the ADC(2) and TDDFT (ωB97X-V) levels for the Chl *d* @ Chl_D1_ model and comparison with the experimental spectrum. (C) Effect of D1-Tyr120 and D1Thr155 on the absorption spectrum computed at the TDDFT level, and comparison to experimental data. The absorption spectrum shows how selectively removing either D1-Tyr120 and D1-Thr155 blueshifts the far-red transition. (D) Structure of the QM region of the tetrameric pigment model, including the P_D1_, P_D2_, Chl_D1_, Pheo_D1_, and surrounding amino acids and water molecules. Hydrogen bonds are shown in dashed orange lines, and hydrogen atoms are omitted for clarity.

For the first-principles treatment of electron correlation effects, we applied the second-order algebraic diagrammatic construction theory, ADC(2), in combination with systematic virtual space reduction (RVS)^19^ and the Laplace transformed (LT) scaled opposite-spin (SOS) treatment.^20-22^ All chlorophyll types showed low *D*_1_ diagnostics in the ADC(2) calculations, indicating a single-reference ground state that provides a reliable basis for the excited state calculations (Table S2). The RVS-LT-SOS-ADC(2) approach allowed us to extend our model to account for the complete reaction center chlorophyll pigment core (6 pigments: 4 chlorophylls and 2 pheophytins) in their protein surroundings, providing a quantitative treatment of electronic excited states,^23^ an approach that has shown promising results for a range of photobiological systems.^22, 24-27^ All QM/MM models were also treated at the LR-TDDFT level in combination with modern range-separated functionals (e.g. ωB97X-V^28^; see *Methods*, Table S3), predicting results highly similar to those from our correlated QM/MM calculations. Our chlorophyll spectra, calculated as an ensemble average over QM/MM molecular dynamics (QM/MM-MD) simulations, reproduce the spectra computed using QM/MM optimized models (Figure S4D), thus indicating that our methodology is robust, despite the computationally highly challenging treatment of electronic excited states in extended multi-pigment models. However, as the absorption maxima of the FRL- and WL-PSII differ by only 0.1 eV and are thus within the error limit of the employed methodology, we rely here also on structural and energetic considerations by probing the influence of the environment on the Chl *d* and *f* substitutions with the RC.

At the QM/MM-ADC(2) level (Figure S4A), when Chl *d* or Chl *f* are modeled at the Chl_D1_ site (with Chl *a* in the other RC pigment locations), we obtain long wavelength transitions at 1.769 eV/701 nm (Chl *d* @ Chl_D1_) and 1.749 eV/709 nm (Chl *f* @ Chl_D1_), respectively. Moreover, when Chl *d* is modeled at the P_D2_ site (Chl *d* @ P_D2_), a prominent absorption feature at 1.703 eV/728 nm is observed, while with Chl *f* at the P_D2_ site (Chl *f* @ P_D2_), an even longer wavelength absorption at 1.606 eV/772 nm arises from the local excitation at P_D2_ (Figure S5). Both models with P_D2_ substitutions are strongly redshifted, but the Chl *d* substitution is closer to the experimental absorption at 1.72 eV/720 nm.^7^

In all four FRL-PSII models as well as in the WL-PSII model, a shorter wavelength transition at 674 nm (1.840 eV) can be observed, corresponding to a local excitation at the P_D1_/P_D2_ pair (Figure S5). This spectral feature arises both at the ADC(2) (Figure S4A) and TDDFT (Figure 1 and Figure S4C) level, suggesting that overall our results are accurate, even when considering the intrinsic error sources of the methods.^23, 29^

Conversely, the TDDFT-spectrum of the pigment clusters with Chl *d* at the Chl_D1_ site aligns well with the experimental spectrum for both the far-red band (709 nm/1.749 eV) and the P_D1_/P_D2_ band at 674 nm (1.840 eV), accurately reproducing of the relative intensities of the Q band (Figure 2, and Table S4, Figure S4A). Taken together, our findings suggest that Chl *d* at the Chl_D1_ site reproduces key optical features of the FRL-PSII, with further energetic and structural implications (see below).

### The effect of the long-wavelength pigment location on the low-energy spectrum

To break down the individual spectral tuning effects, we calculated site energies (Q_y_) by estimating VEEs (S_0_→S_1_) of the pigments, first *without* accounting for the excitonic coupling effect of the surrounding pigments (Table S5 and S6), and then subsequently added the coupling effects arising from interactions with surrounding pigments and the protein (see section below; Figure 3, and Supporting Information, *Extended Methods*).

**Figure 3.**
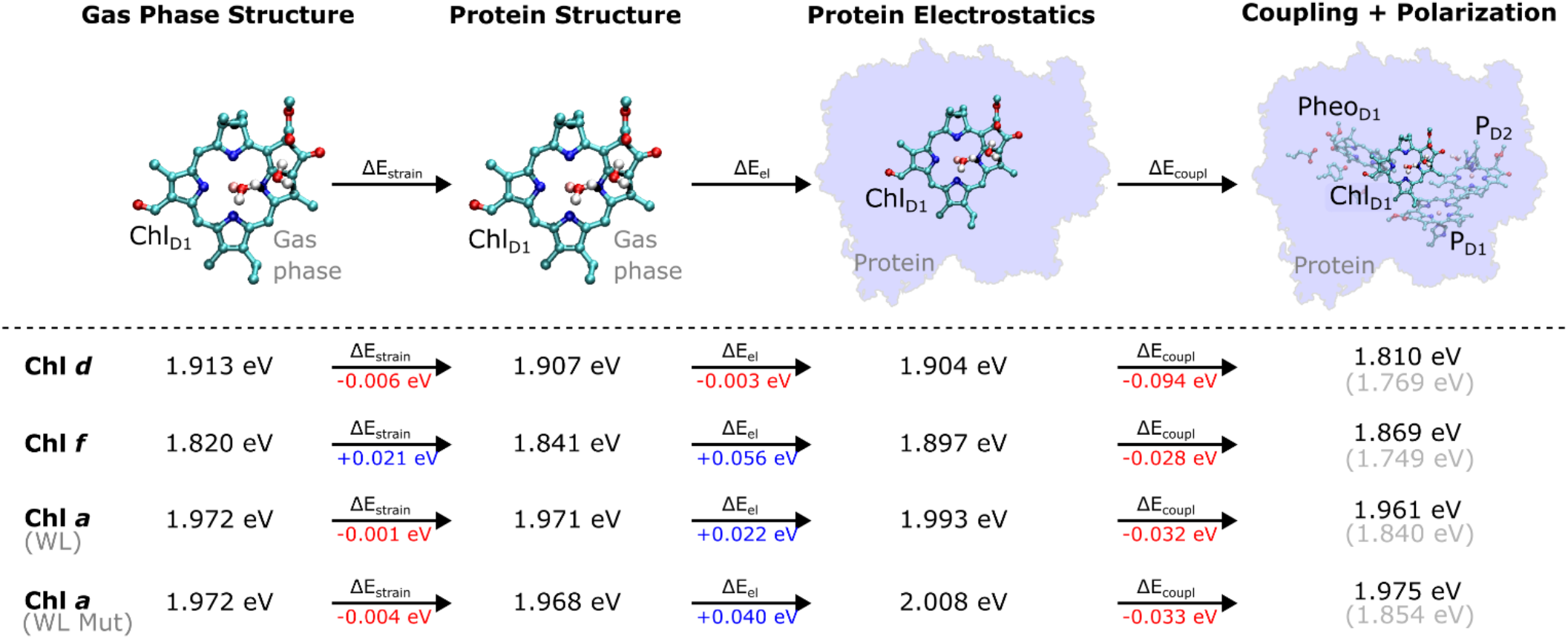
Probing the spectral tuning mechanism of FRL-PSII. The tuning effect was decomposed into electrostatic, strain, and coupling/polarization for the lowest Q_y_ excitation for each of the three pigment types at the Chl_D1_ site. Redshifting values are marked in red, while blueshifting values are marked in blue. Electrostatic effects were estimated by computing shifts in VEEs of QM/MM-optimized pigments from the gasphase to the protein, whilst strain effects were derived from shift of the VEE between the QM/MM-geometry to the gas-phase structure. The grey values in parenthesis refer to the shifted values aligned to the 674 nm peak (see *Methods*). “WL Mut” refers to the WL-PSII model, with F119Y and A154T mutations. All VEEs are derived from ADC(2) calculations, except the “coupling + polarization” contribution, which was calculated at the TDDFT/ ωB97X-V level.

The Chl_D1_ site shows the lowest Q_y_ excitation energies when either Chl *d* (1.904 eV) or Chl *f* (1.897 eV) are modeled at the Chl_D1_ site (Table S5, Table S6). The local excitation on the Chl_D1_ is coupled to a Chl_D1+_/Pheo_D1-_ charge transfer (CT) (Figure S6), consistent with experimental observations^30, 31^ and supporting that the long wavelength pigments could act as a trap for the excitation energy. The differences between the Q_y_ energies of Chl_D1_ and P_D1_/P_D2_ are rather small (0.032 eV for Chl *d*, 0.046 eV for Chl *f*) (Table S5), suggesting that the exciton could be shared amongst multiple sites.

For the WL-PSII model, the lowest Q_y_ excitation comprises the P_D1_/P_D2_ pair together with the Chl_D1_ site (Table S5-S6). In comparison, the site energy of the Chl *a* pigment at Chl_D1_ in the WL-PSII model (1.993 eV) is at shorter wavelengths, as expected, relative to Chl *d* (1.904 eV) and Chl *f* (1.897 eV) (Table S5). The excitonic couplings between P_D1_ and P_D2_ are significantly larger when Chl *d* (97 cm^-1^) or Chl *f* (105 cm^-1^) occupies the Chl^D1^ site, whereas for Chl *a* in the WL-PSII model we obtain an excitonic coupling of 77 cm^-1^ that is slightly lower than the experimental estimates of 80-150 cm^-1^ (see Supporting Information, *Extended Methods*, Table S12).^30-32^

When Chl *d* or Chl *f* occupy the P_D2_ site, the lowest Q_y_ excitation energies are predicted at the P_D1_/P_D2_ pair (1.867 eV for Chl *d*, 1.786 eV for Chl *f*). Relative to the WL-PSII model (1.946 eV) these Q_y_ energies are significantly red shifted. The first excited state (S_1_) of the P_D1_/P_D2_ pair comprises a local excitation on P_D2_ with a small contribution from P_D1_ (Figure S5) that can be considered a “delocalized” state, whilst we obtain stronger excitonic coupling of 180-192 cm^-1^ amongst the Chl *a*/*d* and *a*/*f* pairs at the P_D1_/P_D2_ site.

Overall, the site energies support that the Chl_D1_ substitutions lead to trapping of the excitation energy at the Chl_D1_ site, similar to what is known about the charge separation process in the WL-PSII, whilst the P_D2_ substitutions could result in trapping of the excitation energy at the P_D1_/P_D2_ site and a non-dominant charge transfer pathway.^33, 34^ Taken together, the predicted site energies thus support that the Chl_D1_ substitutions lead to a productive charge separation pathway in the FRL-PSII.

### Color tuning of Chl_D1_ site by the protein-pigment interactions

To understand how the protein environment affects the spectral tuning process, we probed the role of protein-induced macrocyclic ring strain and electrostatic effects in addition to the coupling and polarization effects with the surrounding pigments for the Chl_D1_ site (Figure 3; see *Methods*). The strain effects were calculated based on the spectral shift of an isolated chlorophyll pigment in the gas phase relative to its protein (QM/MM) geometry. We find that the strain effects have a minor redshifting effect on the VEE of Chl *d* (−0.006 eV), whereas it has an overall blueshifting effect on Chl *f* (+0.021 eV) possibly due to the non-preferred binding of the chlorophyll *f* to the Chl_D1_ site (see below).

Interestingly, the protein electrostatics, including hydrogen-bonding effects, also have a redshifting effect for Chl *d* (−0.003 eV), but blueshifts Chl *f* (+0.056 eV) for the Chl_D1_ site, further suggesting that protein electrostatics favors Chl *d* binding. The differences in electrostatic tuning effects can be attributed to the distinct charge density difference upon the photoexcitation (Figure S11) that is favored by the proximity of the C3 formyl group on Chl *d* to the hydrogen-bonding D1-Tyr120/ D1-Thr155 (Figure S7), resulting in an overall redshift for Chl *d*, but no hydrogen-bond and an overall blueshift for Chl *f* (see also Figure 2C, Figure S8). When D1-Tyr120 and D1-Thr155 are modeled into the WL-PSII (replacing D1-Phe119 and D1-Ala154), a blueshift occurs of only 5 nm (0.015 eV) in the Q_y_ band of the Chl *a* at the Chl_D1_ site (Figure 3, Table S7).

Excitonic coupling with surrounding pigments and quantum mechanical polarization effects further redshift the excitation energy (0.032 eV) relative to the monomeric pigment for Chl *d* (Figure 3). Overall, these findings suggest that the intrinsic light-capturing properties of the Chl *d* pigment are enhanced relative to Chl *f* at the Chl_D1_ site by a combination of strain, electrostatics, and coupling/polarization effects, which result in tuning its color and redox properties as the primary electron donor (see below).

### Comparing the functional dynamics of the FRL-PSII and WL-PSII

To explore how the protein dynamics are affected by the FRL substitutions, we studied the FRL-PSII and WL-PSII with Chl *d* or Chl *f* modeled at the Chl_D1_ or P_D2_ sites with atomistic molecular dynamics (MD) simulations (*see* Table S1 for model constructs and analysis). The MD simulations of FRL-PSII with Chl *d* at Chl_D1_ reveal a stable hydrogen-bonding interaction between the formyl group (−CHO) of Chl *d* at the C3 position and both D1-Tyr120 and D1-Thr155 (Figure 4). In the MD trajectory, the D1-Tyr120-formyl interaction is slightly favored over the D1-Thr155-formyl hydrogenbond, whilst our QM/MM-MD simulations favor the latter, possibly due to polarization effects (Figure S10). In contrast, Chl *f* does not form similar hydrogen-bonding interactions at the Chl_D1_ site, as the formyl group occupies the more distant C2 position (Figure 1C, Figure S1C), both rotamers point away from D1Tyr120 throughout the simulations (Figure 4A-C). Similarly, we observe no interactions between D1Tyr120/D1-Thr155 when Chl *a*, which lacks a formyl group at the C3 position, is modeled at the Chl_D1_ site (see also Figure 4A-C).

**Figure 4.**
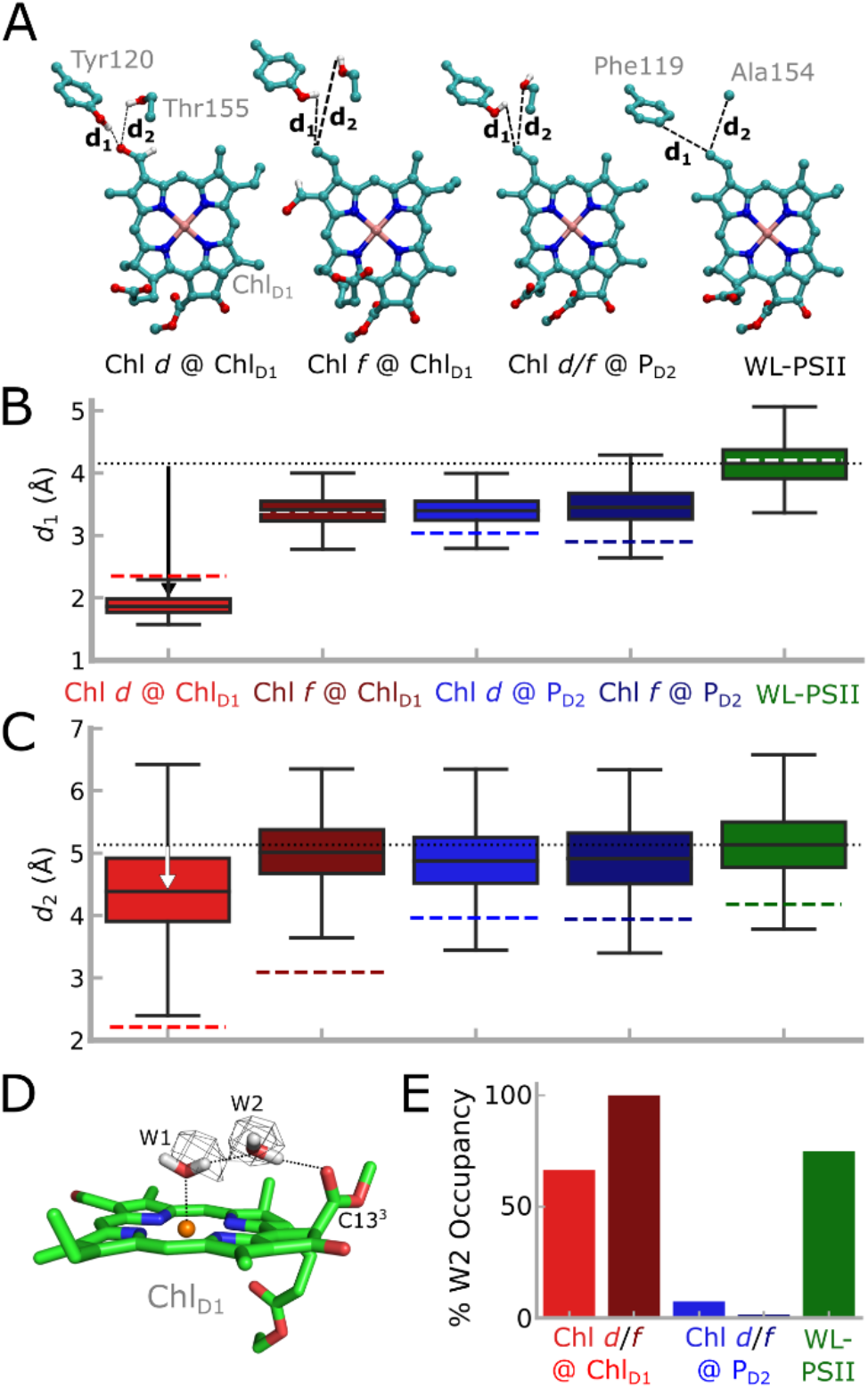
Analysis of MD simulations of the FRL-PSII and WLPSII. (A) Chl_D1_ and its potential hydrogen-bonding partners near the formyl group, namely D1-Tyr120 and D1-Thr155 (D1-Phe119 and D1-Ala154 in the WL-PSII). (B,C) Hydrogen-bonding distance between vinyl (Chl *a*)/formyl (Chl *d*/*f*) group of Chl_D1_ and the (B) D1Tyr120/D1-Phe119 or (C) D1-Thr155/D1-Ala154 (FRL-PSII/ WL-PSII) from the MD simulations. The dotted black line shows the average distance from the WL-PSII MD simulation as a reference. MD distances are shown as a box plot, while QM/MM optimized distances are indicated as colored dashed lines. Boxes indicate the second and third quartiles of the distribution. (D) Water molecules around Chl_D1_. The water molecule W2 is bound between W1 and the C13_3_ carbonyl group of Chl_D1_. The cryo-EM density of the FRL-PSII (PDB ID: 7SA3, EMD-24943)_12_ is consistent with the water positions obtained from MD simulations (see also main text). (E) Occupancy of the W2 position in each of the MD models. In the Chl *d/f* @ P_D2_ simulations W2 escapes the binding pocket.

The P_D2_ pigment is stabilized through hydrogen-bonding interaction to D2-Tyr191 via the C13^2^ group in the FRL-PSII model (D2-Trp191 in WL-PSII), as well as stacking interactions with P_D1_. To this end, no hydrogen-bonding interactions are seen between the protein and the formyl groups Chl *d* or *f* when they occupy the P_D2_ site (Figure S10A). The formyl group of Chl *d*, when in the P_D2_ location, is highly dynamic in the MD simulation, sampling multiple rotameric states. In contrast, the formyl group of Chl *f* remains in a single rotameric conformation during the MD simulations (Figure S10B).

In addition to these local interactions, we observe differences in the dynamics of the hydrogen-bonding network around the Chl_D1_, especially for a specific water molecule, W2. In the MD simulations W2 forms a hydrogen-bonding link (67% occupancy) between W1, the water that is the axial ligand to Mg^2+^ of Chl_D1_, and its carbonyl (C13^2^-COOCH_3_) substituent (Figure 4D-E). W2 is structurally conserved at the Chl_D1_ site, and its density is well-resolved in both the WL-PSII and FRLPSII.^5, 12, 35^ Modifications of the coordinating water molecule and its hydrogen-bonding network can affect both optical and redox properties of the chlorophyll (Table S3, Figure S10).^36-38^

In contrast, for our MD simulations of Chl *d* and Chl *f* at the P_D2_ site, W2 near Chl_D1_ escapes from the pigment pocket, possibly due to steric effects, repositioning D2Leu205 and the propionate group of Chl_D1_ that could affect the local hydration (Figure 4). These changes are further supported by the smaller sidechain of D1Leu173 in the FRL-PSII relative to D1-Met172 in the WL-PSII (Figure S10D).

Our MD simulations suggest that both Chl *d* and Chl *f* at Chl_D1_ lead to subtle structural changes within the RC. We observe a small increase in the P_D1_/P_D2_ inter-pigment distance (3.65 Å average, *edge-to-edge*) with Chl *d* at the Chl_D1_ site relative to the WL-PSII (3.53 Å) (Figure S10A) that could influence the electronic coupling (Figure S13), charge separation, and electron-hole localization following the charge-separation process. For Chl *f* at the Chl_D1_ site, the P_D1_/P_D2_ inter-pigment distance increases to 3.73 Å, most likely due to steric effects.

In contrast, our MD simulations with Chl *d* and Chl *f* at P_D2_, revealed only minor perturbation in the inter-pigment spacing of P_D1_/P_D2_ (3.55 Å for both Chl *a*/*f* and Chl *a*/*d* composition). Nevertheless, our calculations suggest that the presence of the long wavelength chlorophyll would affect both the excitonic coupling (Figure S13) and redox properties of the pigments (see below; Table S9).

### Redox tuning and charge transfer kinetics in the FRLPSII

To probe the binding affinities of the different chlorophylls to the RC, we estimated the binding energies using a PBSA/MM-model based on the MD trajectories (see Supporting Information, *Extended Methods*). Our analysis suggests that Chl *d* at the Chl_D1_ site has a stronger binding affinity relative to Chl *f* or Chl *a* at this position (Figure S12B,D). This effect can be attributed to hydrogen-bonding from D1-Tyr120 or D1Thr155, stabilizing Chl *d* relative to the other pigments. For the P_D2_ site, Chl *a* is better stabilized in comparison to Chl *d/f* (Figure S12C). This likely reflects steric effects from the formyl group, but also the preference for water over histidine as the axial ligand in Chl *f*, though not in Chl *d* (Figure S12B). Moreover, our binding calculations suggest that the P_D2_ site prefers Chl *a* (Figure S12B).

To probe how the Chl *d* substitution at the Chl_D1_ site affects the charge separation in the FRL-PSII, we estimated redox potentials based on Poisson-Boltzmann electrostatic calculations coupled with Monte Carlo sampling (PBE/MC), and probed the charge separation kinetics using a kinetic master equation model based on electron transfer theory (see Supporting Information, *Extended Methods*).

Our PBE/MC calculations suggested that the oxidation potential of the Chl_D1_ site increases by 145 mV in the FRL-PSII model with Chl *d* and by 198 mV with Chl *f* relative to the WL-PSII model (Figure S16). Following the excitation forming the P_720*_ state at 1.72 eV, the charge separation in the FRL-PSII is thus expected to result in a Chl_D1+_/Pheo_D1-_ state at 1.57 eV with Chl *d* at the Chl_D1_ site, computed based on the shifts in redox potentials relative the WL-PSII. To this end, the PBE/MC calculations suggest that the Chl *d* modifications at Chl_D1_, stabilize the charge-separated states due to the hydrogen-bonding effects to surrounding residues by *ca*. 0.15 eV relative to the WL-PSII system (Figure 5A), and could thus compensate for the 0.10 eV lower photon energy in the FRL-PSII, while maintaining a significant driving force needed for the water oxidation and quinone reduction.

Interestingly, our PBE/MC models suggest that with Chl *f* at Chl_D1_, the Chl_D1+_Pheocharge separated-state becomes more stabilized possibly due to its intrinsically higher oxidation potential relative to Chl *d* and *a* (Figure 5A, Figure S16). The shifted redox properties result in less driving force to stabilize the subsequent electron transfer steps and could favor re-combination and harmful singlet oxygen production, as it is energetically close to the Mn_ox_/Q_B-_ state (1.25 eV).

**Figure 5.**
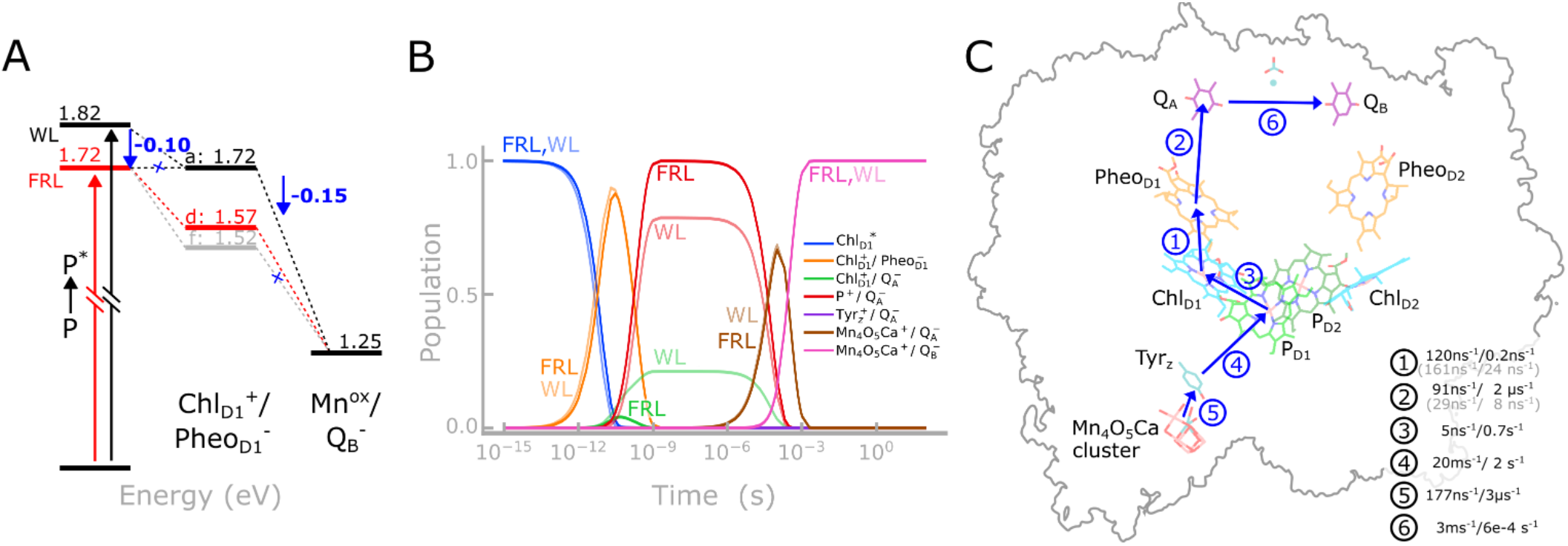
Probing the charge separation mechanism of FRL-PSII. (A) Effect of the chlorophyll substitution at the Chl_D1_ site on the charge separation energetics based on redox calculations and experimental data. (B) CT kinetics and state populations along the Chl_D1+_Pheo_D1-_-mediated pathway for the FRL-PSII (Chl *d* @ Chl_D1_; bright colors) and WL-PSII (transparent colors). (C) Kinetics of charge transfer: Light-driven CT at Chl^D1^ leads to a Chl_D1+_/Pheo_D1-_ state, whilst the subsequent electron transfer leads to reduction of Q_B_ and oxidation of the Mn_4_O_5_Ca cluster. The arrows indicate the order of events predicted by our kinetic model. The charge transfer rates reported correspond to the FRL system (with Chl *d* at the Chl_D1_ site), whilst rates that are unique for the WL-PSII model, are shown in grey. See Figure S15, S16 and Table S10 and Supporting Information, *Extended Methods* for detailed models.

Our kinetic models based on the determined energetics (Figure 5B, 5C, Figure S15), predict that the Chl_D1+_Pheo--mediated charge separation pathway is fast, with a halftime reaching Q_B_ reduction/Mn oxidation within *ca*. 300 µs (Figure 5B) (cf. Refs. 39, 40), which is rate-limited by the slow Q_A_→ Q_B_ reaction.^41^ The shifted redox properties at Chl_D1_ thus support a charge separation kinetics comparable to that in the WL-PSII, despite the lower photon energy and shifted redox properties.

## DISCUSSION

Our multiscale calculations and prediction of excitation energies showed that the chemical substitution of the Chl_D1_ site by Chl *d* provides both a structural and electronic model of the FRL-PSII that is consistent with experimental optical spectra^7^ and cryo-EM structural analysis^12^ (see also *Introduction*). At this site, Chl *d* is orientated appropriately to accept a hydrogen-bond from the far-red specific, conserved residues D1Tyr120 and D1-Thr155, that tune the absorption, the redox potentials, and increase its affinity to this site.

When Chl *f* occupies the Chl_D1_ site, our predictions also showed redshifted features that could be consistent with the experimental spectra. However, Chl *f* in the Chl_D1_ site is not capable of binding to the far-red specific, conserved hydrogen-bonding residues, and the presence of which resulted in a small blueshift of the Chl *f* Q_y_ absorption and a lowered binding constant.

When Chl *d* and Chl *f* were modeled at the P_D2_ site, it led to a large redshift of the longest wavelength absorption significantly beyond the experimentally observed range (Figure 2A, Figure S4A). Moreover, these models did not show similar structural stabilization as for the chemical substitution at the Chl_D1_ site, as there were no obvious conserved changes in the FRL-PSII that could accommodate, either sterically or by the provision of hydrogen-bonds, the additional formyl groups present on Chl *d* or *f*. In addition, the presence of Chl *f* at the P_D2_ site perturbs the structure of the pigment cluster resulting in the loss of the W2 water that plays a role in the hydrogen-bond network associated with the W1 water that coordinates the Chl_D1_. Changes in this hydrogen-bonding network can result in spectroscopic and functional changes making the PSII susceptible to photodamage.^37, 38^

ssTaken together, our combined findings thus indicate that the *Chroococcidiopsis thermalis* FRL-PSII has a long wavelength chlorophyll at the Chl_D1_ site, with Chl *d* reproducing key electronic and structural features, and support initial arguments for the Chl_D1_ location for the long wavelength pigments (see *Introduction*).^7, 9, 12^

Our excited-state calculations and analysis of transition orbitals (Figure S6, S7) showed that a long wavelength chlorophyll at the Chl_D1_ site gave rise to a low-energy Q_y_ excitation that could act as the primary donor and result in a Chl_D1+_/Pheo_D1-_ state (Figure 5A). This primary radical pair state transfers the cation radical (the electron hole) to P_D1_/P_D2_ pair and transfers the electron to Q_A_, creating a transient charge-separated (P_D1_P_D2_)_+_/Q_A-_ state, with a *t*_1/2_ of 0.2 ns (Figure 5B), as suggested by our kinetic model, and with subsequent electron transfer steps forming Tyr_z·_(H_+_)Q_A-_ and eventually Mn_ox_Q_B-_ as the final state (Figure 5C). Interestingly, the chemical substitution of Chl_D1_ compensates for the lower photon energy by the tuned redox properties that could further favor the charge transfer towards the Q_B_ sites (Figure 5A). In this regard, recent 2D-electronic spectroscopy (2D-ES) studies of the FRLPSII^42^ found experimental support for a similar (P_D1_P_D2_)_+_/Chl_D1-_ mediated pathway, although this charge separation was assumed to be initiated from a state in which Pheo_D1_ is pre-reduced. Our kinetic models indicate that Chl *a* at the Chl_D1_ site lacks the driving force necessary for initial charge separation in the FRLPSII, while for Chl *f* at this site the Chl_D1+_/Pheo_D1-_ state is energetically further stabilized, and could potentially promote electron back transfer and charge recombination, which in turn leads to the formation of triplet chlorophyll and the production of harmful singlet oxygen. Chl *d* is unique among the pigments as it supports reasonable charge separation energetics at the Chl_D1_ site, and facilitates fast charge transfer kinetics, which are comparable to the WL-PSII according to our kinetic model. In this regard, our electrostatic calculations, suggest that the charge-separated Chl_D1+_/Pheo_D1-_ pair is stabilized by *ca*. 0.15 eV to compensate for the *ca*. 0.1 eV lower photon energy. Although our current models reveal distinct differences between the FRL- and WL-PSII, elucidating the exact mechanistic details of the charge transfer kinetics will require further detailed structure-based electron transfer calculations.

In contrast, when either Chl *d* or *f* occupies the P_D2_ site, our calculations suggest that the excitation energy is likely to be trapped within the RC (Figure S15, Table S5, S6). The energy transfer to Chl_D1_, which is the putative site of primary charge separation in the WL-PSII (see above; and Refs. 31, 33, 43-45), would thus involve an energetically uphill and kinetically slow excitation transfer process. While modification of the pigment at the P_D2_ site would indeed expand the spectral range of PSII (Figure 2A, Figure S4A), we speculate that the putative (P_D1_P_D2_)_+_/Chl_D1-_ state with Chl *f* at P_D2_ would result in a charge recombination.

Our calculations suggest that the specific FRL-PSII surroundings around Chl_D1_ support the redshifting effect of Chl *d* that could enhance its intrinsic light-capturing properties at the far-red-light limit, while Chl *f* at the same position led to blueshifting effects on the intrinsic low-energy Q_y_ state. We could link these spectral shifts to protein-induced electrostatic and hydrogen-bonding effects, leading to an energetic preference of the Chl *d* at the Chl_D1_ site. This effect arises specifically via the FRL-PSII specific D1-Tyr120 and D1Thr155 residues that form hydrogen-bonding interactions with Chl *d*. In summary, considering both the pigment-protein interactions at the Chl_D1_ site, spectral tuning effects, as well as possible bioenergetic advantages in the charge separation process (Figure 5), our calculations strongly support that Chl *d* at the Chl_D1_ site enables the FRL-PSII to capture low energy photons beyond the far-red light limit.

## CONCLUSION

We have combined here multiscale QM/MM models with atomistic molecular dynamics simulations and excited state calculations to elucidate molecular details underlying far-red light adaptations of Photosystem II. Based on our multiscale calculations and analysis of experimental data, we found that modifications of both pigment and protein are critical for enhancing the far-red light absorbing properties of FRL-PSII, which achieves spectral tuning by combined electrostatic and polarization/excitonic coupling effects. Taken together, our work strongly supports the presence of a Chl *d* pigment at the Chl_D1_ site within the RC, leading to a model yielding low site energies that can initiate the primary charge separation reaction and drive water oxidation. Our study provides a basis for further structure-based assignment of color-tuning mechanisms and bioenergetic principles of oxygenic photosynthesis at the far-red light limit.

## MATERIALS AND METHODS

Models of the WL-PSII (PDB ID: 3WU2)_5_ and FRL-PSII (with Chl *a*/*d*/*f* at P_D2_ or Chl_D1_) were embedded in a POPC/water/ion simulation box with 100 mM NaCl, and studied using atomistic MD simulations with the CHARMM36m force field_46, 47_ together with in-house parameters of the co-factors. The MD simulations were propagated for 300 ns in duplicates with a 2 fs integration timestep at *T*=310 K and *p*=1 bar using NAMD2.14/NAMD3._48, 49_ The trajectories were analyzed with VMD._50_ Optical spectra were computed at the QM/MM level, with 1-6 pigments (100-600 atoms) modeled at the TDDFT or LT-SOS-ADC(2)/def2-TZVP_18, 51_level and the MM region at the CHARMM36m level. The QM/MM simulations were performed using the CHARMM/ TURBOMOLE interface._52-54_Electronic couplings were calculated using the fragment-excitation difference (FED) approach_55_, as implemented in QChem v. 5.3._56_ Redox properties were estimated by combining quantum chemically computed reduction/oxidation free energies with classical solvation free energies modeled at a linearized Poisson-Boltzmann level, as implemented in APBS._57_ See further details of the multiscale simulations in the Supporting Information, *Extended Methods*.

## ASSOCIATED CONTENT

### Supporting Information

Extended methods; figures showing ADC(2) and TDDFT spectra, natural transition orbitals, binding free energies, excitonic coupling; site energies; reduction and oxidation potentials; benchmarking; computational details. This material is available free of charge via the Internet at http://pubs.acs.org.

## ACKNOWLEDGMENT

This work was supported by the Knut and Alice Wallenberg Foundation (V.R.I.K. grant: 2019.0251), the Swedish Research Council (V.R.I.K.), the Royal Society Wolfson Research Merit Award (A.W.R.) and Royal Society Research Professorship 2024 (AWR) and Biotechnology and Biological Sciences Research Council (BBSRC) grants (A.W.R. BB/R001383/1, BB/V002015/1). Moreover, the TU Munich-Imperial College Collaboration Fund provided initial support to V.R.I.K. and A.W.R. We also acknowledge the TUM Global Incentive Fund, funded under the Excellence Strategy of the Federal Government and the Länder. V.R.I.K. acknowledges also the DFG for their support within the Mercator Fellow Program to SFB1078, while A.S. acknowledges the EMBO Long-Term Fellowship (ALTF 952-2022) for their support. For computational resources, we thank the Leibniz Rechenzentrum (LRZ, project:pr83ro) and the National Academic Infrastructure for Supercomputing in Sweden (NAISS 2024/1-28, NAISS 2023/1-31), as well as the National Infrastructure for Computing (SNIC 2022/1-29 and SNIC2021/1-40) at the Center for High Performance Computing (PDC), partially funded by the Swedish Research Council through a grant agreement no. 2018-05973.

---

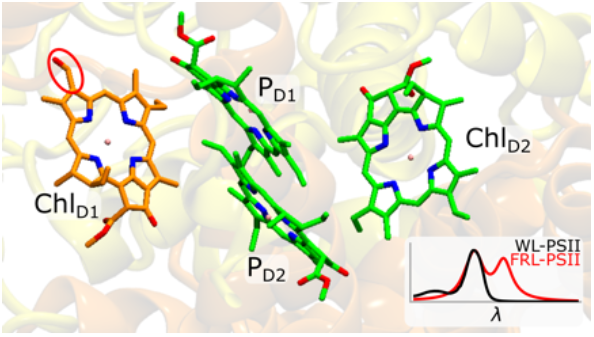

