## Supporting Information for "Modified Chlorophyll Pigment at Chl_D1_ Tunes Photosystem II Beyond the Red-Light Limit"

### Table of Contents

#### Extended Methods

*Atomistic MD models*
*DFT models*
*QM/MM calculations*
*Micro-iterative QM/MM optimizations of extended QM models*
*Excited state calculations*
*Poisson-Boltzmann Electrostatics (PBE) coupled to Monte-Carlo (MC) sampling of chlorophyll redox states.*
*Calculation of reduction/oxidation potentials of model compounds.*
*Kinetic Models*

#### List of Figures

**Figure S1.** Structure of the FRL-PSII and WL-PSII.
**Figure S2.** Sequence comparison of the FRL-PSII and WL-PSII.
**Figure S3.** Computational models.
**Figure S4.** Comparison of calculated absorption spectra.
**Figure S5.** Natural transitions orbitals for $P_{D1}/P_{D2}$.
**Figure S6.** Natural transitions orbitals for $Chl_{D1}/Pheo_{D1}$.
**Figure S7.** Natural transitions orbitals for $Pheo_{D1}$-$Chl_{D1}$-$P_{D1}$-$P_{D2}$.
**Figure S8.** Effect of protein charge distribution on the absorption spectra.
**Figure S9.** Structure and different conformations of Chl *a*, Chl *d*, and Chl *f*.
**Figure S10.** Conformational dynamics of Chl *d* / *f*.
**Figure S11.** Density difference upon photo-excitation for different chlorophyll pigments.
**Figure S12.** Binding free energies of chlorophyll clusters in different PSII isoforms.
**Figure S13.** Excitonic couplings within $P_{D1}/P_{D2}$.
**Figure S14.** The micro-iterative optimization scheme of the QM/MM models.
**Figure S15**. Comparison of simulated charge transfer kinetics in the WL-PSII and FRL-PSII.
**Figure S16**. Calculated oxidation potentials.

#### List of Tables

**Table S1.** List of QM/MM models and MD simulations.
**Table S2.** $D_1$ diagnostics for RVS-LT-SOS-ADC(2) calculations of different chlorophylls types.
**Table S3.** Benchmarking VEEs for Chl *a*.
**Table S4.** Calculated excitation energies of the FRL-PSII and WL-PSII.
**Table S5.** Summary of VEEs for the FRL-PSII models from ADC(2) calculations.
**Table S6.** Summary of VEEs for the FRL-PSII models from TDDFT calculations.
**Table S7.** VEEs for different pigments at the $Chl_{D1}$ position.
**Table S8.** Distribution of charge/spin upon oxidation of $P_{D1}/P_{D2}$.
**Table S9.** Calculated and experimental reduction and oxidation potentials used as model compounds for Chl *a*, Chl *d*, and Chl *f*.
**Table S10.** Parameters used for modeling the charge transfer kinetics.
**Table S11**. Experimental oxidation and reduction potentials of redox active pigments and cofactors in WL-PSII.
**Table S12**. Excitonic couplings of central pigments in the WL-PSII and FRL-PSII.
**Table S13.** RVS screening for LT-SOS-ADC(2) calculations of Chl *d* at the $Chl_{D1}$ position.

#### SI References

#### Extended Methods

**Atomistic MD models.** The MD simulation models were constructed based on the x-ray structure of the WL-PSII from *Thermosynechococcus vulcanus* (PDB ID: 3WU2)[1] with the D1, D2, CP43 and PsbH subunits modified for the FRL-PSII model first using MODELLER[2] (see sequence analysis Figure S2), followed by optimization of the cofactors with the CHARMM36m force field using in-house parameters [3, 4]. Four Chl *a* pigments were changed to Chl *f* and one Chl *a* pigment to Chl *d*: the antenna chlorophylls C507, C608, and C612 were replaced with Chl *f*, while C508 and $Chl_{D1}$/$P_{D2}$ were changed to either Chl *f* or Chl *d* (Table S1, Figure S1A). This resulted in four different FRL-PSII simulation models (Chl *d* @ $Chl_{D1}$, Chl *f* @ $Chl_{D1}$, Chl *d* @ $P_{D2}$ and Chl *f* @ $P_{D2}$) in addition to our WL-PSII model with all Chl *a*. Structures of Chl *d* and Chl *f* were constructed based on Chl *a*, with the chlorophyll pigments linked to either one axial water molecule or a histidine residue. The PSII complexes were embedded in a 1-palmitoyl-2-oleoyl-*sn*-glycero-3-phosphocholine (POPC) membrane,[5] solvated with TIP3 water molecules, and neutralized with 100 mM NaCl ions. The final system contained approximately 535,000 atoms (Figure S3C). MD simulations (3 μs in total) were performed using the CHARMM36m force field,[6] with all cofactor parameterized through in-house DFT calculations (at the B3LYP-D3/def2-TZVP level).[3, 4] Atomic partial charges of the cofactors in different redox and spin states, were derived based on the RESP[7] procedure, and force constants based on the molecular Hessian or from model compounds. The pigments were parametrized in the neutral, oxidized, and reduced states, including Chl *a*/*d*/*f* with and without axial (water/His) ligands, and $Pheo_{D1}$. The simulations were carried out in the $S_2Y_z$ state of the $Mn_4O_5Ca$ cluster in PSII [3]. The MD simulations were performed at $T$=310 K and $p$=1 atm, a 2 fs integration timestep, and computing the long-range electrostatics using the particle mesh Ewald (PME) method (grid size of 1 Å). The models were initially relaxed by a 4 ns relaxation run with harmonic restraints of 1 kcal $mol^{-1}$ $Å^{-1}$, followed 54 ns equilibration without restraints, and 300 ns production runs in duplicates. The simulations were carried out using NAMD2.14/NAMD3,[8, 9] and the trajectories were analyzed using VMD.[10]

**DFT models.** DFT models of isolated Chl *a*, Chl *d*, Chl *f*, and Pheo *a* were constructed based on the atomic structure of Chl *a* or Pheo *a* obtained from the WL-PSII structure (PDB ID: 3WU2)[1]. The quantum chemical models comprised 71-73 atoms. The chlorophyll tails were cut between the $C17^1$ and $C17^2$ atoms, and terminal carbon atoms were saturated with hydrogen atoms. The structures were geometry optimized at the B3LYP-D3/def2-SVP/def2-TZVP ($Mg^{2+}$) level of theory,[11-14] followed by TDDFT and RVS-LT-SOS-ADC(2) calculations[15] with def2-TZVP basis sets. For the TDDFT treatment, we tested different functionals (B3LYP, CAM-B3LYP[16], CAMh-B3LYP[17], LRC-ωPBEh[18], ωB97X-V[19], ωB97M-V[20] (Figure S4C). All calculations were performed using TURBOMOLE v. 7.5[21] and VMD[10] used for analysis.

**QM/MM calculations.** Hybrid QM/MM calculations (Figure S3B) were performed based on the WL-PSII and FRL-PSII models (see above) with Chl *d*/Chl *f* in either the $Chl_{D1}$ or $P_{D2}$ for the FRL-PSII, resulting in a total of five different models (Table S1). Geometry optimizations were performed with around 66,000 MM atoms and a QM region comprised two RC chlorophyll, $P_{D1}$ and $P_{D2}$, together with their coordinating histidine residues (D2-His197 and D1-His199/D1-His198 for FRL/WL) and six coordinating water molecules, as well as the neighboring chlorophylls ($Chl_{D1}$, $Chl_{D2}$), D1-Phe119/D1-Tyr120 (WL/FRL), and D1-Ala154/D1-Thr155 (WL/FRL). The two pheophytins ($Pheo_{D1}$, $Pheo_{D2}$) were also included together with D1-Gln130/D1-Glu131 (WL/FRL). The QM region comprised 510-512 atoms (Figure S3A), which were sequentially optimized in sub-steps (see Figure S14). Link atoms were introduced between $C_\alpha$ and $C_\beta$ atoms, or for chlorophyll tails between $C17^1$ and $C17^2$ atoms. The QM/MM systems were optimized using the adopted basis Newton-Raphson algorithm at the B3LYP-D3/def2-SVP / MM level of theory, while QM/MM-MD simulations were propagated using a 1 fs timestep at $T$=310 K. All QM/MM calculations were performed using the CHARMM/TURBOMOLE interface.[6, 21, 22]

**Micro-iterative QM/MM optimizations of extended QM models.** The QM region of the QM/MM calculations comprised 510-512 atoms, which were stepwise optimized using a micro-iterative approach (see Figure S14). In the first iteration, the $P_{D1}$ and $P_{D2}$ pigments together with their histidine ligands were optimized in an initial iteration. In the second step, $Chl_{D1}$ and $Chl_{D2}$ together with three surrounding water molecules each and two amino acids interacting with $Chl_{D1}$ (D1-Tyr120/D1-Thr155 for the FRL-PSII or D1-Phe119/D1-Ala154 for the WL-PSII), were added to the QM/MM model, increasing the size of the QM region to four pigments. In the third optimization step, QM/MM models comprising $Pheo_{D1}$ and its coordinating D1-Glu131 (FRL-PSII)/D1-Gln130 (WL-PSII) were optimized, followed by a fourth optimization round, where $Chl_{D1}$ together with its surrounding residues (see above) was merged with the $Pheo_{D1}$ QM/MM model, and re-optimized. In step 5, the second pheophytin ($Pheo_{D2}$) was optimized, followed by the final optimization step, where the four central pigments ($Chl_{D1}$, $P_{D1}$, $P_{D2}$ and $Chl_{D2}$) were re-optimized (same QM region as in step 2). The resulting QM/MM structures, obtained after steps 1-6, were used for calculations of spectra.

**Excited state calculations.** Excited state calculations were carried out using the QM/MM optimized structures of the FRL-PSII and WL-PSII models with Chl *d* and Chl *f* modeled either at the $Chl_{D1}$ or $P_{D2}$ position. Models of single chlorophyll pigments were used for the estimation of site energies (Table S5, Table S6), while tetrameric models consisting of $P_{D1}$-$P_{D2}$-$Chl_{D1}$-$Pheo_{D1}$ were used for the calculation of electronic spectra. These models included in addition to the four central pigments, D2-His197 and D1-His199 (ligands of $P_{D1}$ and $P_{D2}$), D1-Tyr120 and D1-Thr155, three water molecules for $Chl_{D1}$, and D1-Glu131 (Figure 1B). The protein environment was modeled with electrostatic embedding, with the MM system described by point charges at the CHARMM36m force field level.

To probe the color tuning mechanisms in the FRL-PSII, Chl *d*/*f* was modeled at either the $P_{D2}$ or the $Chl_{D1}$ positions. Excited state properties of the individual pigments were computed with the RVS-LT-SOS-ADC(2)/def2-TZVP level of theory,[12, 23] with virtual orbitals 40 eV[24] above the highest occupied molecular orbital (HOMO) frozen. The systematic RVS error was accounted for by extrapolating all values to the frozen-core limit, yielding $\Delta VEE$=-0.090 eV for Chl *a*/*d* models and $\Delta\Delta VEE$=-0.085 eV for Chl *f*, based on isolated chlorophyll models (Table S13). The absorption spectrum for the $P_{D1}$-$P_{D2}$-$Chl_{D1}$-$Pheo_{D1}$ QM/MM model was calculated at the RVS-LT-SOS-ADC(2)/def2-TZVP level based on computations on the full spectrum from the $P_{D1}$-$P_{D2}$ and $Chl_{D1}$-$Pheo_{D1}$ models.

LR-TDDFT calculations were performed at the ωB97X-V/def2-TZVP level for individual pigments and assemblies, as described above. For benchmarking purposes, we also performed TDDFT computations with the B3LYP, CAM-B3LYP, CAMh-B3LYP, LRC-ωPBEh, ωB97X-V, and ωB97M-V DFT functionals. Depending on the pigment composition, we computed a total of 14 and 16 excited states at the ADC(2) and TDDFT levels, respectively. Vertical excitation energies (VEEs) and corresponding oscillator strengths were used to compute the full spectrum using Lorentzian broadening. In this regard, the spectra were visualized by convoluting the determined peaks with a Lorentzian function of width 12 nm. The computed spectrum has uniformly shifted and aligned with respect to the principle 674 nm peak to aid the comparison.

The chlorophyll spectrum was also computed as an ensemble average based on 32.2 ps QM/MM-MD sampling from selected MD snapshots (23 x 1.4 ps distributed equally over a cumulative 600 ns MD trajectory). The resulting spectra was obtained from 913 individual VEEs obtained at the TDDFT/MM level (Figure S4D). The QM/MM-MD simulations were propagated using a 1 fs timestep at $T$=310 K.

The electronic coupling between the pigments was computed using the fragment-excitation difference (FED) scheme implemented in QChem v.5.3[25] at the TDDFT (ωB97X-V/def2-TZVP) level of theory. The protein environment was included using the point charges in electronic coupling calculations. The QM/MM optimized geometries were utilized to assess the influence of inter-pigment distances on electronic coupling parameter.

**Poisson-Boltzmann Electrostatics (PBE) coupled to Monte-Carlo (MC) sampling of chlorophyll redox states.** Binding energies and redox potentials of Chl *a*/*d*/*f* in PSII were estimated based on the continuum electrostatic calculations by solving the linearized Poisson-Boltzmann equation (PBE) as implemented in the adaptive Poisson-Boltzmann Solver (APBS) [26] and titrating residues and cofactors with Karlsberg+ [27-29]. Partial charges of the cofactors in different oxidation states were modeled as before,[3] and Chl *d*/*f* and Pheo parameters, derived as described above (see DFT models section). To this end, we estimated oxidation potentials for $P_{D2}$, and both reduction and oxidation potentials for $Chl_{D1}$ for model compounds, based on DFT calculations of reference redox potentials (see below; Table S9). Redox potentials and $pK_a$ values were computed by modelling the protein as force field point changes embedded in a polarizable ε=4 surroundings and ε=80 for water (see ref. 30), with the cofactors modelled in different oxidation states. The model was truncated to include the lipids and water molecules within 5 Å of the protein (D1, D2, CP43, CP47 and PsbO). The atoms were modelled using van der Waals radii (based on the CHARMM36m force field), whilst the aqueous surroundings were defined based on a surface exclusion area calculation, with a water probe radius of 1.4 Å. The electrostatic energies were computed by numerical solution of the linearized PBE with a 100 mM ionic strength (KCl), as implemented in APBS,[26] whilst MC sampling was used to find the lowest energy state. An automatized PBE/MC routine, implemented in Karlsberg+,[27-29] was used to optimize hydrogen positions in the most probable protonation/redox states, using five iterations/cycles. Cofactor binding energies were computed using a molecular mechanics/ Poisson–Boltzmann continuum area solvation model (MM/PBSA) based on a thermodynamic cycle and by estimating solvation and electrostatic interactions of the chromophore within the protein environment.[31] The calculated reduction and oxidation potentials and their shifts (Figure S16), together with the experimental reference data (Table S10) were used for modeling the electron/charge transfer kinetics (Main text, Figure 5).

**Calculation of reduction/oxidation potentials of model compounds.** DFT models of pheophytin, as well as water- and histidine-coordinated Chl *a*, Chl *d*, and Chl *f* were constructed based on the crystal structure from the WL-PSII (PDB ID: 3WU2) (Figure S9).[1] The models, comprising 71-103 atoms, were truncated between the $C17^1$ and $C17^2$ atoms, and the terminal carbon atoms were saturated with hydrogen atoms. The structures for neutral, anionic, and cationic species were geometry optimized at the B3LYP-D3/def2-SVP/def2-TZVP ($Mg^{2+}$) level.[11-13] Reduction and oxidation free energies, $\Delta G_{red/ox}$, were calculated at $T$= 298 K based on electronic energies,[32, 33] computed at the B3LYP-D3/def2-TZVP level, and zero-point energies, thermal vibrational corrections, and entropic contribution estimated at the (B3LYP-D3/def2-SVP/def2-TZVP ($Mg^{2+}$) level according to,

$$\Delta G_{red/ox} = (E_{red}-E_{ox}) + (ZPE_{red} - ZPE_{ox}) + [E^{vib}{}_{red}(T) - E^{vib}{}_{ox}(T)] - T\,(S_{red} - S_{ox}) + \Delta\Delta G^{red\text{-}ox}{}_{solvation}$$

with the vibrational corrections, computed from the vibrational energies ($\varepsilon_i$) was obtained from the molecular Hessian at $T$= 298 K,

$$E^{vib}{}_{red/ox}(T) = -\,k_B T \log\left(\prod_i \left[\,1 / (1-\exp(-\varepsilon_i/k_B T))\right]\right)$$

The solvation free energy difference ($\Delta\Delta G^{red\text{-}ox}{}_{solvation}$) between the reduced and oxidized species were estimated using **solvate** module of MEAD [34], where the solute was described with atomic partial charges obtained by the RESP [7] method, and the solvent is represented by an homogeneous dielectric with ◎=80 for water and ◎=37.5 for acetonitrile (AN). To this end, a solvation radius of $r$=1.42 Å was used for water and $r$=2.23 Å for AN, with atomic radii obtained from Ref. 33. Electronic energies and vibrational contributions were calculated using TURBOMOLE v. 7.5.[21]

**Kinetic Models.** The electron transfer (eT) kinetics in the FRL-PSII and WL-PSII were modeled based on Marcus theory, with rates derived from microscopic calculations (see below) and experiments.[35] To this end, the eT rates were computed by considering all possible electron transfer pathways between donor (D) and acceptor (A) atom pairs, and weighting each pathway by the respective coupling elements based on protein packing densities along the given pathway.[36, 37]

$$\log k_{eT,i\to j} = \frac{1}{N_i N_j}\sum_{a=1}^{N_i}\sum_{b=1}^{N_j}\left(13 - (1.2 - 0.8\rho_{ab})(R_{ab}\text{Å}^{-1} - 3.6) - 3.1\frac{\left(\Delta G_{i\to j} + \lambda_{i\to j}\right)^2}{\lambda_{i\to j}}\right),$$

where $\rho_{ab}$ are the pair-weighted packing density, $R_{ab}$ are the distance between all D-A pairs, and $N_i$ and $N_j$ denote the number of atoms in the donor and acceptor selection, respectively. The definition above, differs from the Moser-Dutton formulation, where rates are computed based on the shortest "*edge-to-edge* pathway" (cf. 37). The driving force for the eT, $\Delta G_{i\to j}$ was estimated based on experimental $E_m$ values for the WL-PSII, but shifted by the electrostatic interactions between the D-A pair. In this regard, the interaction was modeled based on a Coulombic interaction, based on explicit molecular calculations with atomic point charges between the electron and hole obtained from DFT-calculations, and using an $\varepsilon$=10. Shifts in the $\Delta G_{i\to j}$ for the FRL models were estimated based on PBC/MC calculations (Figure S16), whereas *r*eorganization energies ($\lambda_{i\to j}$) of cofactors were calculated as,

$$\lambda_{ox} = E_{ox}(r_{opt,red}) - E_{red}(r_{opt,red})$$

and

$$\lambda_{red} = E_{red}(r_{opt,ox}) - E_{ox}(r_{opt,ox}).$$

$$\lambda_{tot} = \lambda_{ox}^{D} + \lambda_{red}^{\mathrm{A}}$$

where $E_{ox}$ and $E_{red}$ refer to electronic states of the cofactors, and $r_{opt,red}$ and $r_{opt,ox}$ are the optimized geometries in the reduced and oxidized states, respectively. The electron transfers kinetics (Figure 5) was modeled by numerically integrating the master equation,

$$\frac{dp_i}{dt} = \sum_j k_{j\to i}p_j - \sum_i k_{i\to j}p_i$$

using the Livermore Solver for Ordinary Differential Equations (LSODA) integrator as implemented in COPASI 4.37.[35]

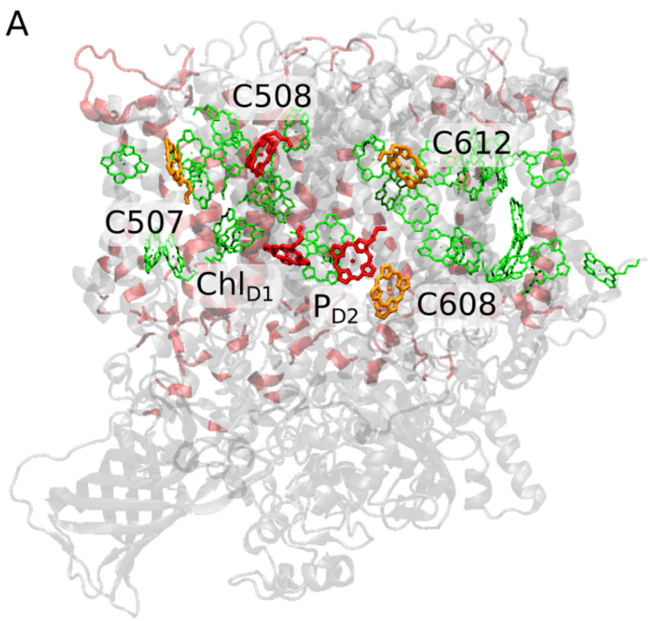


**Figure S1. Structure of the FRL-PSII and WL-PSII.** (A) Structure of the FRL-PSII. Changes in the FRL sequence in comparison to the WL-PSII are highlighted in red. Chlorophyll pigments are shown in green; the three putative Chl *f* molecules in orange; and the three Chl *d* candidates in red, whereas one of them was modelled as Chl *d* and the other two as Chl *f* depending on the model (*see* Table S1 for details on the models). (B) Comparison of the Chl$_{D1}$ surrounding (showing hydrogen-bonding residues) for the FRL-PSII (*left*) and WL-PSII (*right*).

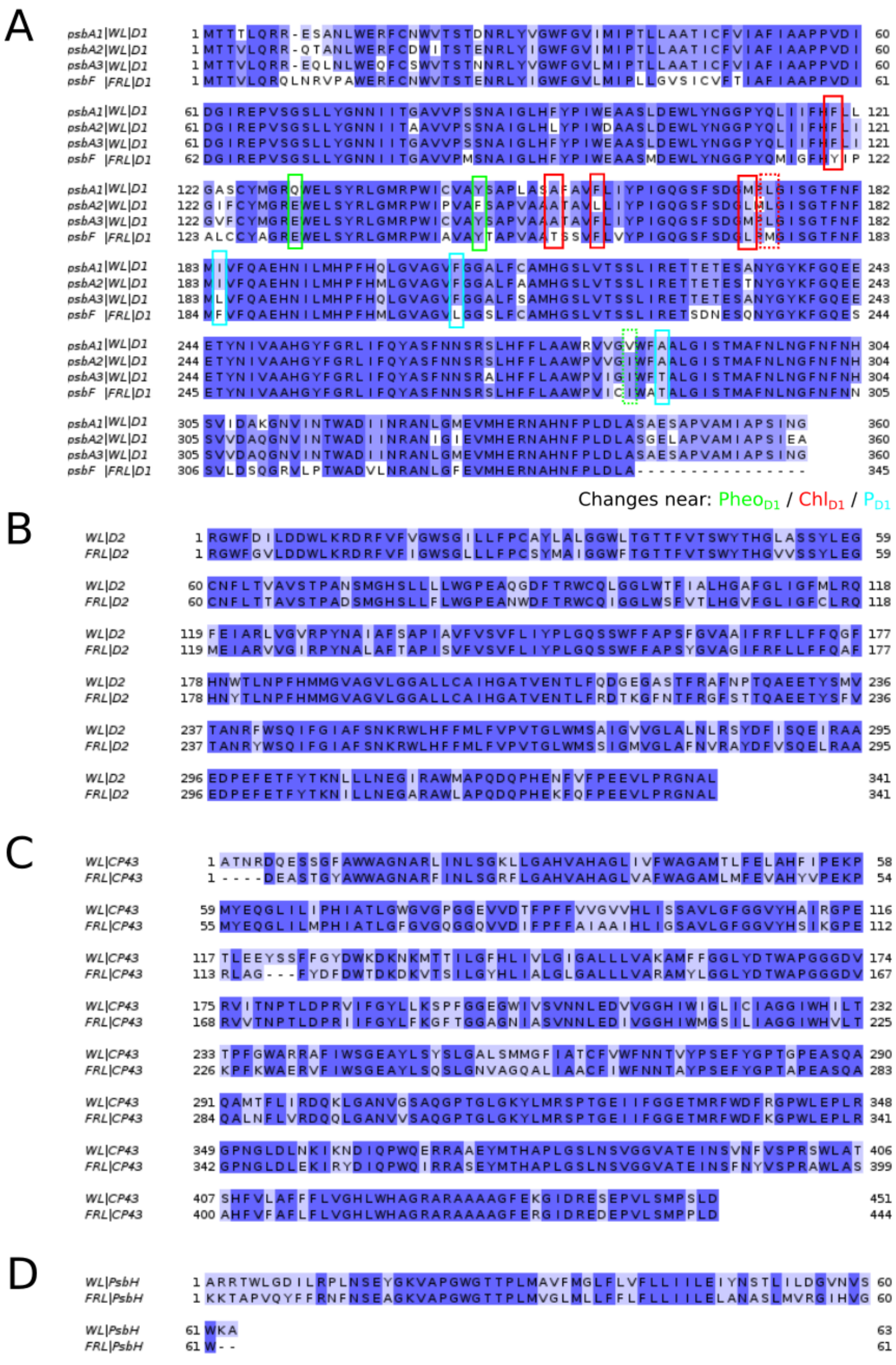

**Figure S2. Sequence comparison of the FRL-PSII and WL-PSII.** Subunits (A) D1, (B) D2, (C) CP43, and (D) PsbH are shown. The sequence identities of the FRL-PSII and WL-PSII are 83%, 84%, 78%, and 60% respectively. All remaining subunits (not shown), of the FRL-PSII models are taken from the WL-PSII. (A) Subunit D1 is encoded by the *psbF* gene in the FRL-PSII, while *psbA1* (3WU2)[1], *psbA2* (7YQ2)[38] and *psbA3* (7YQ7)[38] encode for the WL-PSII, (B)-(D) Changes in the sequence around the reaction core: 3/4/2 single residue substitutions in D1 are observed in the proximity of $P_{D1}$/$Chl_{D1}$/$Pheo_{D1}$, whilst the region around the OEC is fully conserved between the isoforms.

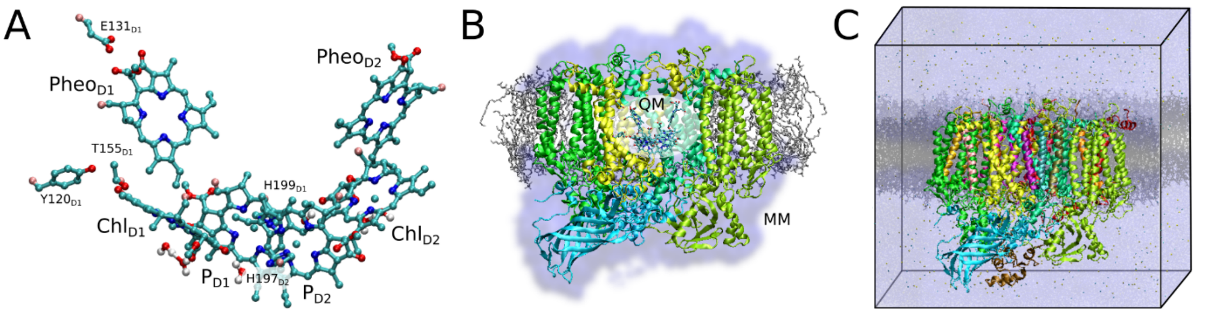


**Figure S3. Computational models.** (A) QM region of the QM/MM models, involving 6 pigments (600 atoms), and surrounding residues / bound water molecules for the FRL-PSII (see Figure S1 for differences to WL-PSII). This QM region was minimized in micro-iterative steps (for technical details, see *Extended Methods*, and Figure S14). (B) QM/MM model of PSII, illustrating the QM region and the surrounding protein system (MM region with 66,000 atoms). The MM region was trimmed to include six center subunits (D1, D2, CP43, CP47, PsbO and PsbV), their embedded cofactors, as well as surrounding lipids (within 5 Å of the QM/MM model). (C) Atomistic membrane-bound model of the (WL/FRL-) PSII used in the molecular dynamics (MD) simulations.

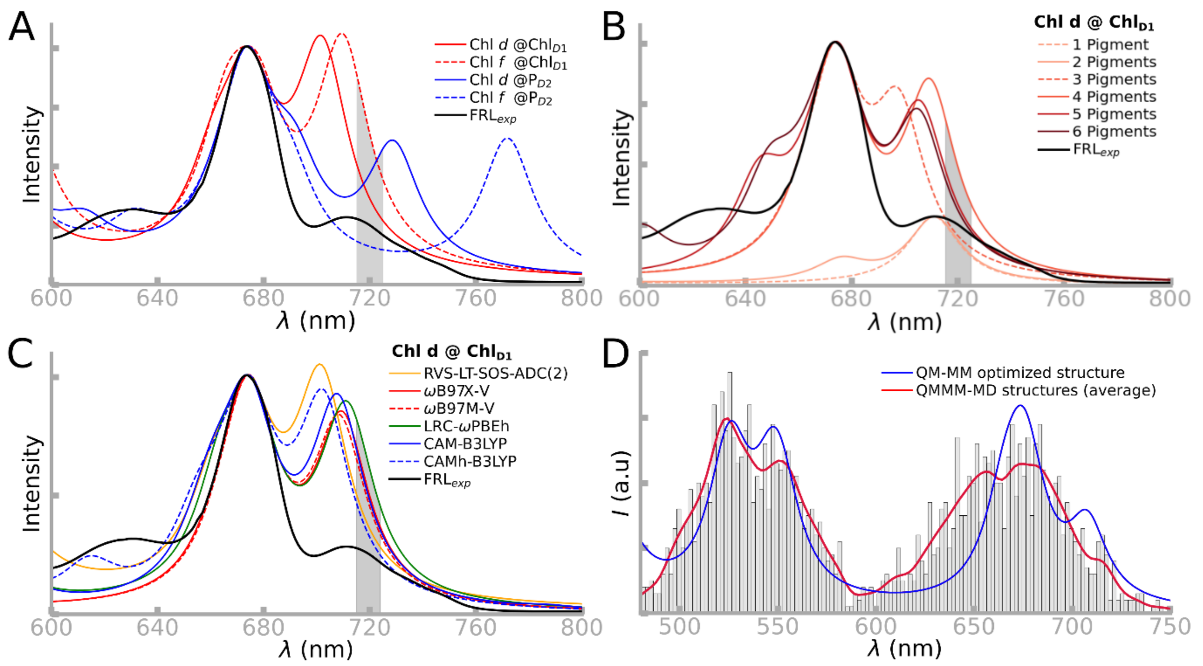


**Figure S4. Comparison of calculated absorption spectra**. (A) RVS-LT-SOS-ADC(2) spectra for the tetramer assembly for Chl *d*/*f* at $Chl_{D1}$ (in red) or $P_{D2}$ (in blue). While the Chl *d*/*f* on $Chl_{D1}$ spectra are in good agreement with the experimental FRL-PSII spectrum, the $P_{D2}$ substitutions show a second peak around 728 nm (Chl *d*) or 772 nm (Chl *f*), not present in the experimental spectrum. (B) Effect of the selection of pigments (at the TDDFT level). "1 Pigment" QM region: $Chl_{D1}$; "2 pigments" QM region: $Chl_{D1}$-$Pheo_{D1}$; "3 pigments" QM region: $P_{D1}$-$Chl_{D1}$-$Pheo_{D1}$; "4 pigments" QM region: $Chl_{D1}$-$Pheo_{D1}$-$P_{D1}$-$P_{D2}$; "5 pigments" QM region: $Chl_{D1}$-$Pheo_{D1}$-$P_{D1}$-$P_{D2}$-$Chl_{D2}$ ; "6 pigments" QM region: $Chl_{D1}$-$Pheo_{D1}$-$P_{D1}$-$P_{D2}$-$Chl_{D2}$-$Pheo_{D2}$. Each of the QM region also includes key amino residues (see *Extended Methods*). All computed spectra show similar features, independent of the exact pigment model used, and supporting that the models are robust. The analysis in the main text is focused on the "4 pigment" model. (C) Effect of the choice of DFT functionals on the absorption spectrum (tetramer model of Chl *d* @ $Chl_{D1}$). LRC-ωPBEh captures the position of the 674 nm and 711-725 nm peak, while ωB97X-V provides an overall good fit. (D) Comparison of the spectra obtained by fitting a single TDDFT spectra from a QM/MM minimized structure (Lorentzian broadening), with the spectra obtained from the ensemble average over QM/MM-MD simulations (see *Extended Methods*).

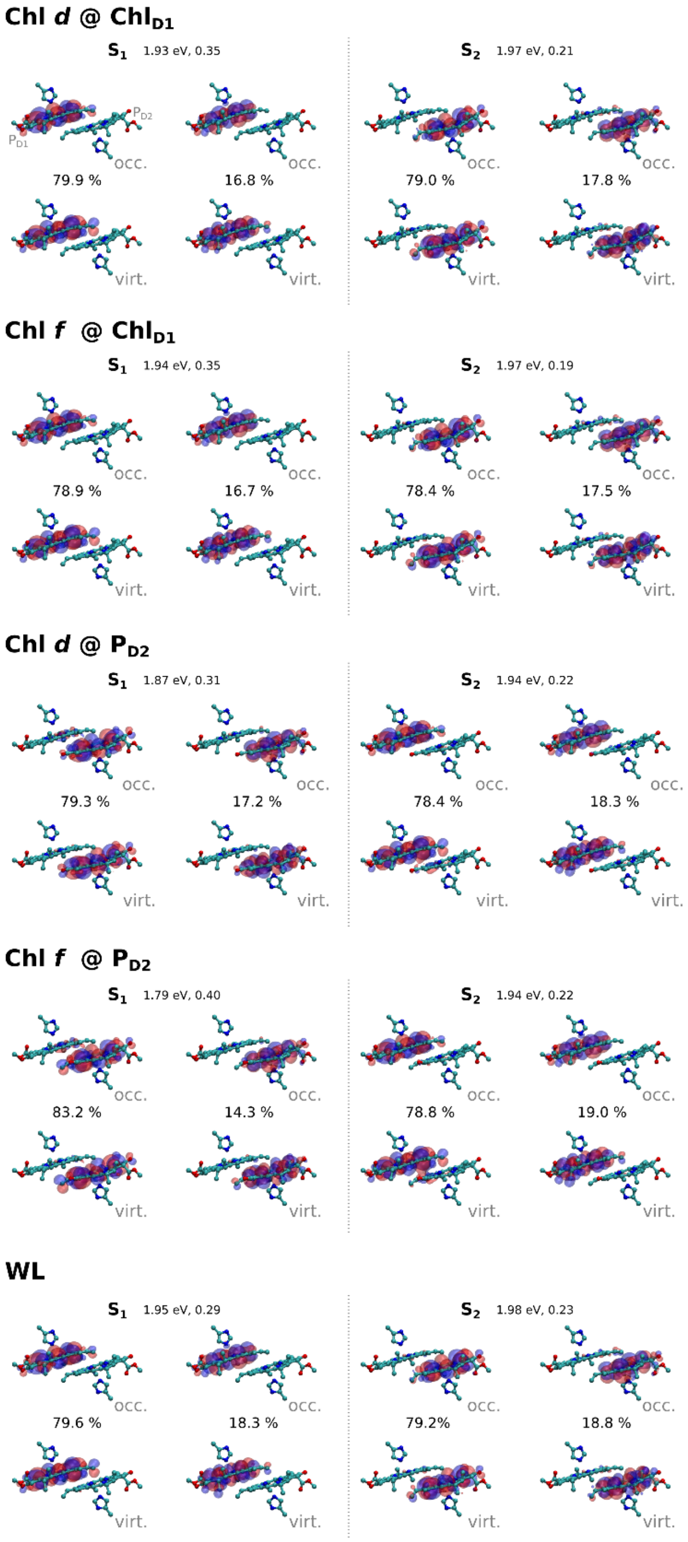


**Figure S5. Natural transitions orbitals for $P_{D1}/P_{D2}$.** The figure shows natural transition orbitals (NTO) for $S_1$ and $S_2$ of the $P_{D1}/P_{D2}$ pair, calculated at the RVS-LT-SOS-ADC(2) level. Both occupied and virtual NTOs are shown along with their respective weights, VEEs, and oscillator strengths. The low-lying states of the $P_{D1}/P_{D2}$ pair comprise local excitations. Positive NTOs are shown in blue, negative in red, with a threshold of ±0.01 e/Å$^3$.

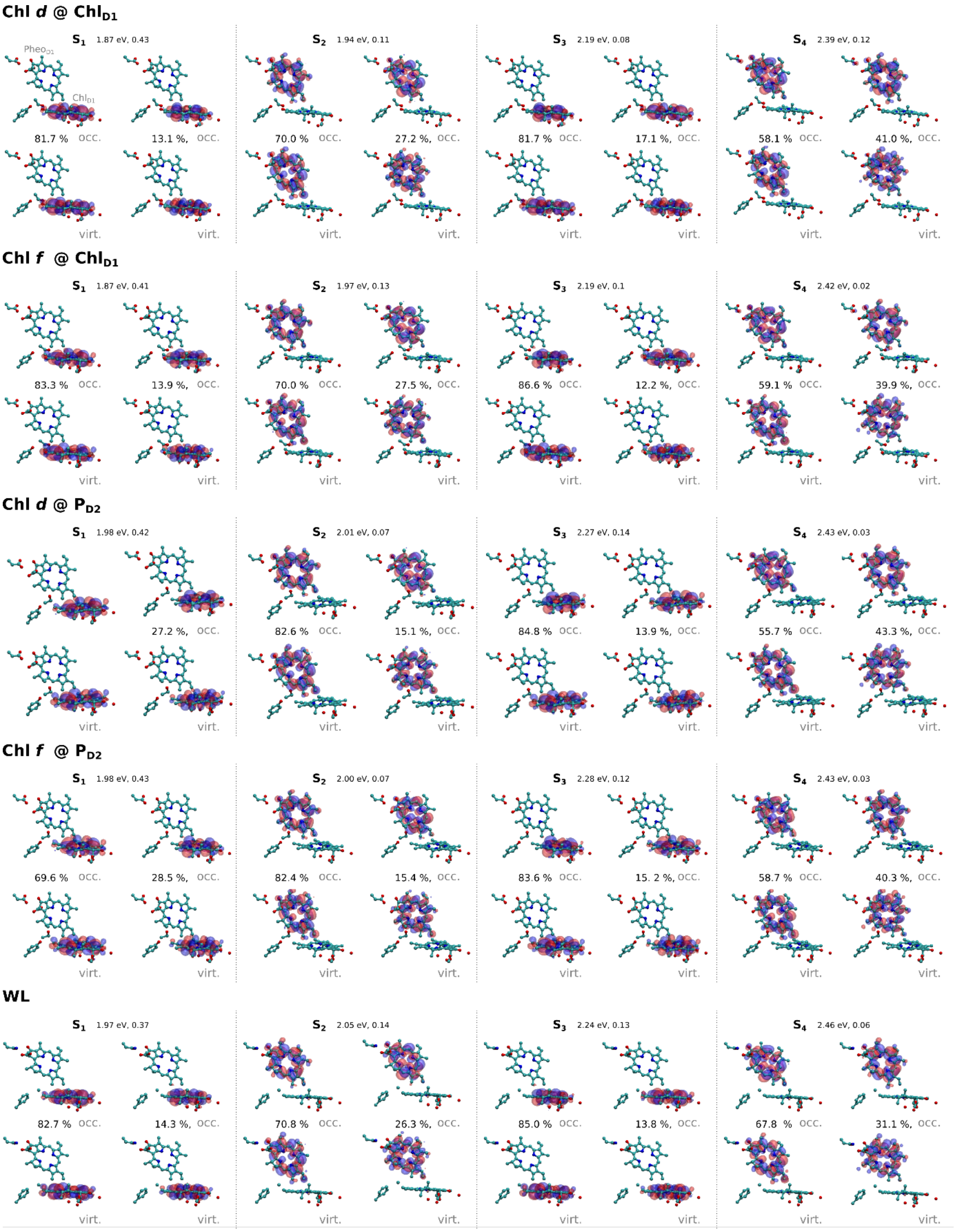


**Figure S6. Natural transitions orbitals for $Chl_{D1}$/$Pheo_{D1}$.** The figure shows natural transition orbitals (NTO) for $S_1$ - $S_4$ of the $Chl_{D1}$/$Pheo_{D1}$ pair, calculated at the RVS-LT-SOS-ADC(2) level. Both occupied and virtual NTOs are shown along with their respective weights, VEEs, and oscillator strengths. The low-lying states of the $Chl_{D1}$-$Pheo_{D1}$ comprise local excitations, mainly the $Q_y$ and $Q_x$ states of the $Chl_{D1}$ and $Pheo_{D1}$. Positive NTOs are shown in blue, negative in red, with a threshold of ±0.01 e/Å$^3$.

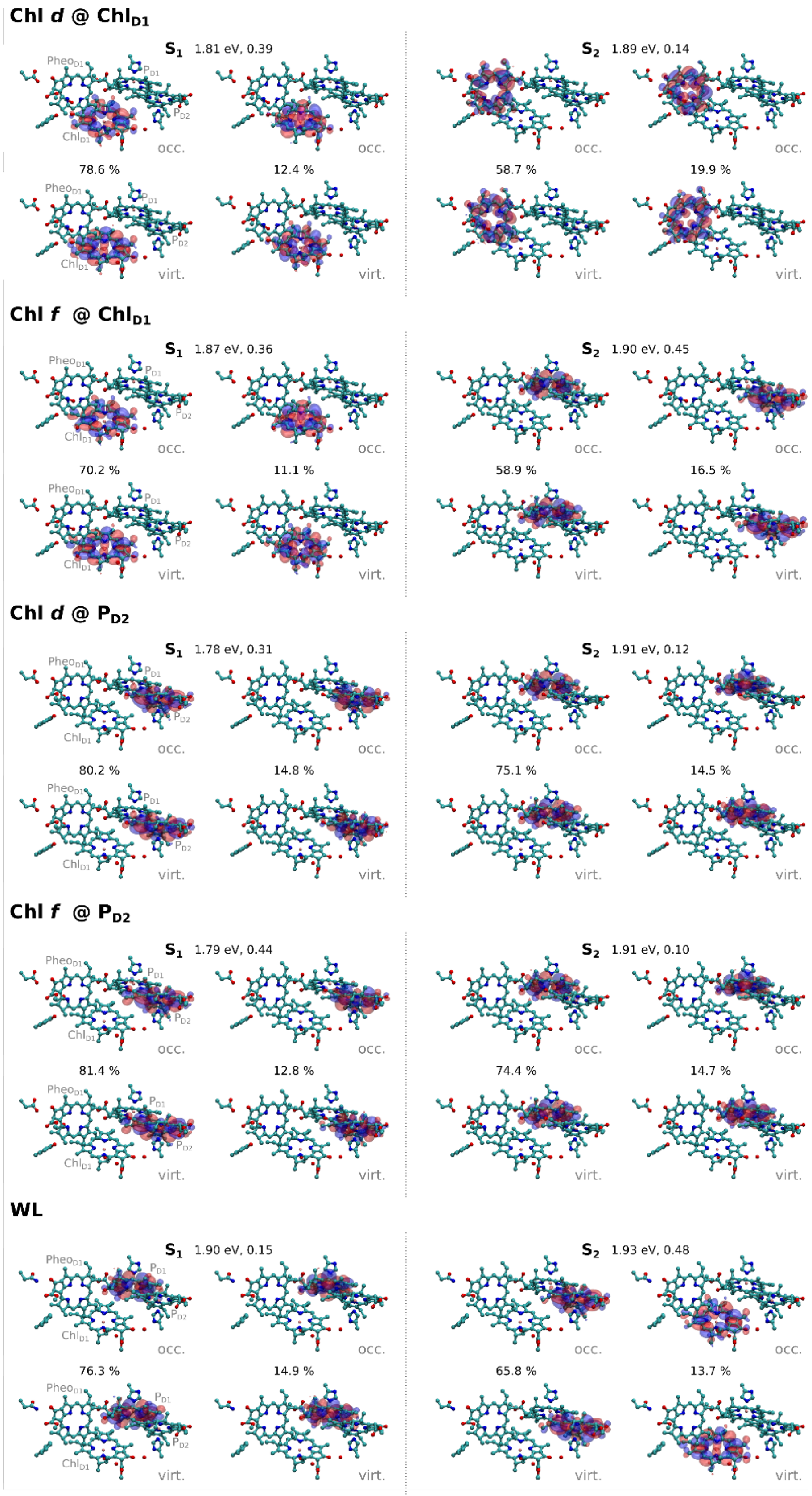


**Figure S7. Natural transitions orbitals for Pheo$_{D1}$-Chl$_{D1}$-P$_{D1}$-P$_{D2}$.** The figure shows natural transition orbitals (NTO) for the Pheo$_{D1}$-Chl$_{D1}$-P$_{D1}$-P$_{D2}$ tetramer for S$_1$-S$_4$, calculated at the ωB97X-V/def2-TZVP level. Both occupied and virtual NTOs are shown along with their respective weights, VEEs, and oscillator strengths. The NTOs are shown at an isosurface value of ±0.01 e/Å$^3$ (positive - blue; negative - red).

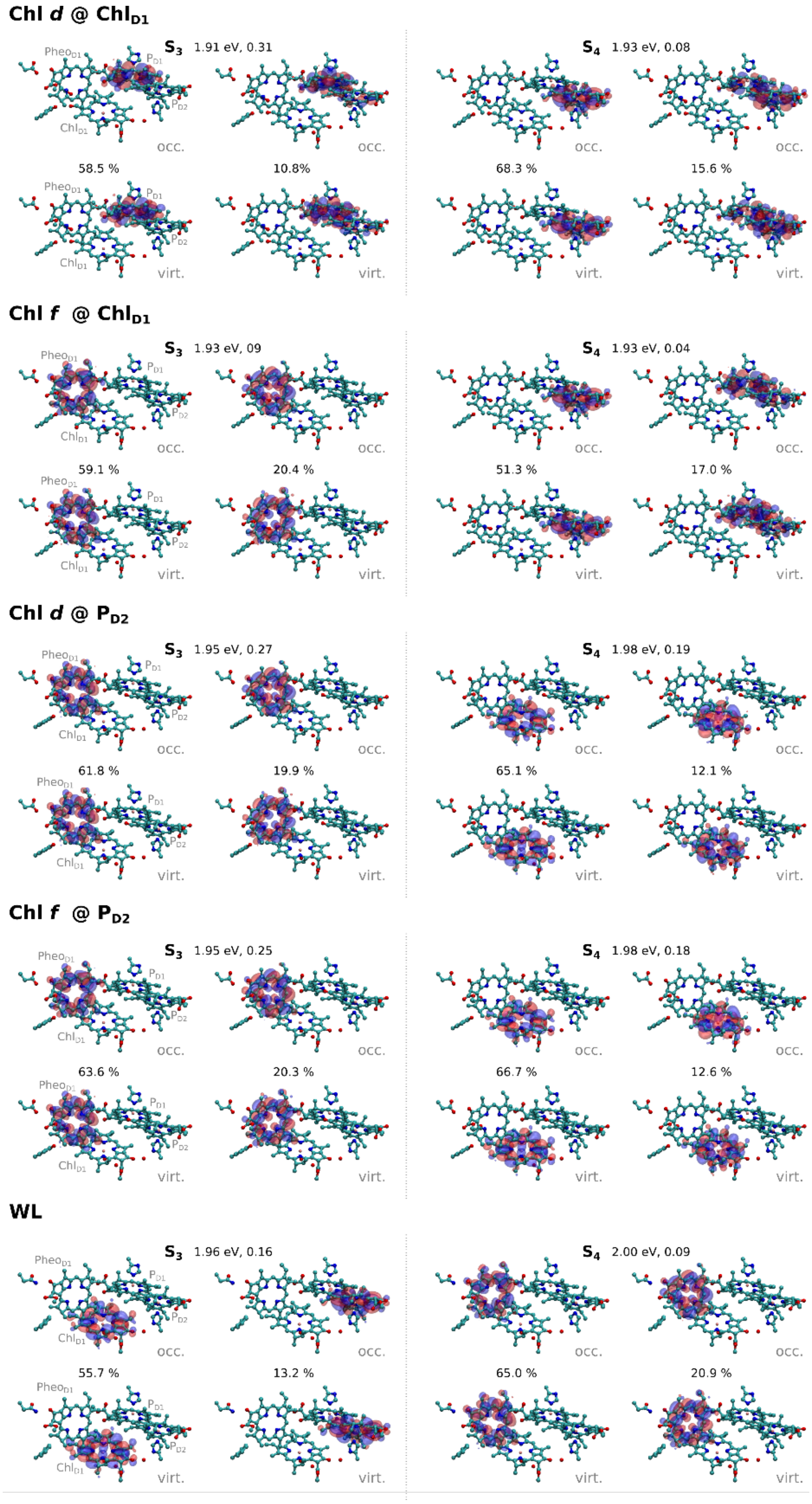


**Figure S7. Natural transitions orbitals for $Pheo_{D1}$-$Chl_{D1}$-$P_{D1}$-$P_{D2}$** (***contd.***). The figure shows natural transition orbitals (NTO) for the $Pheo_{D1}$-$Chl_{D1}$-$P_{D1}$-$P_{D2}$ tetramer for $S_1$-$S_4$, calculated at the ωB97X-V/def2-TZVP level. Both occupied and virtual NTOs are shown along with their respective weights, VEEs, and oscillator strengths. The NTOs are shown at an isosurface value of ±0.01 e/Å$^3$ (positive - blue; negative - red).

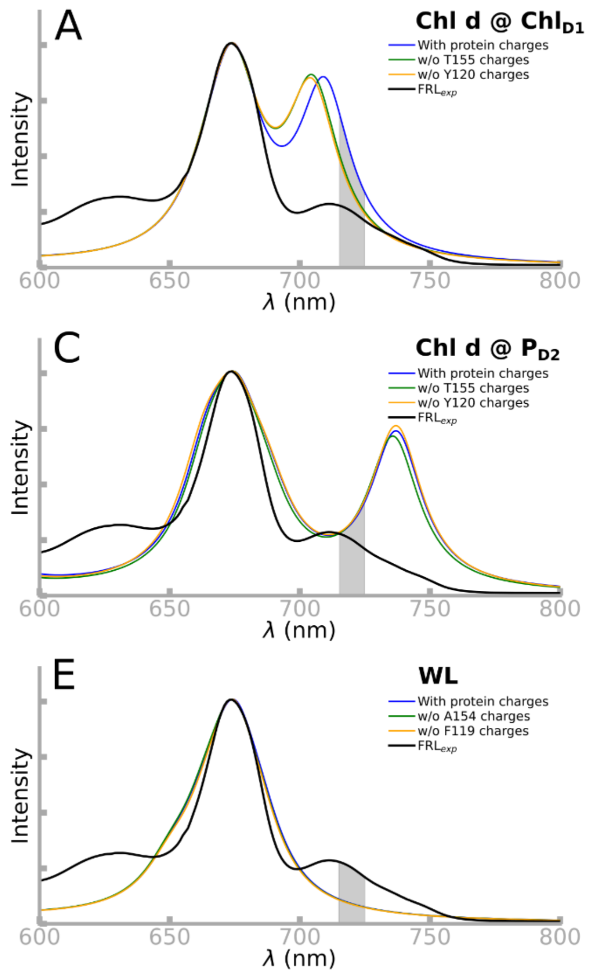


**Figure S8. Effect of protein charge distribution on the absorption spectra.** The calculated absorption spectra show the effect of selectively removing either D1-Tyr120 or D1-Thr155, leading to a blueshifted far-red peak. The calculations were performed on a tetrameric QM model at the TDDFT (ωB97X-V/def2-TZVP)/MM level. (A)-(E) Spectra calculated for the five different QM/MM models (see Table S1), with the model indicated in the top right corner of each panel.

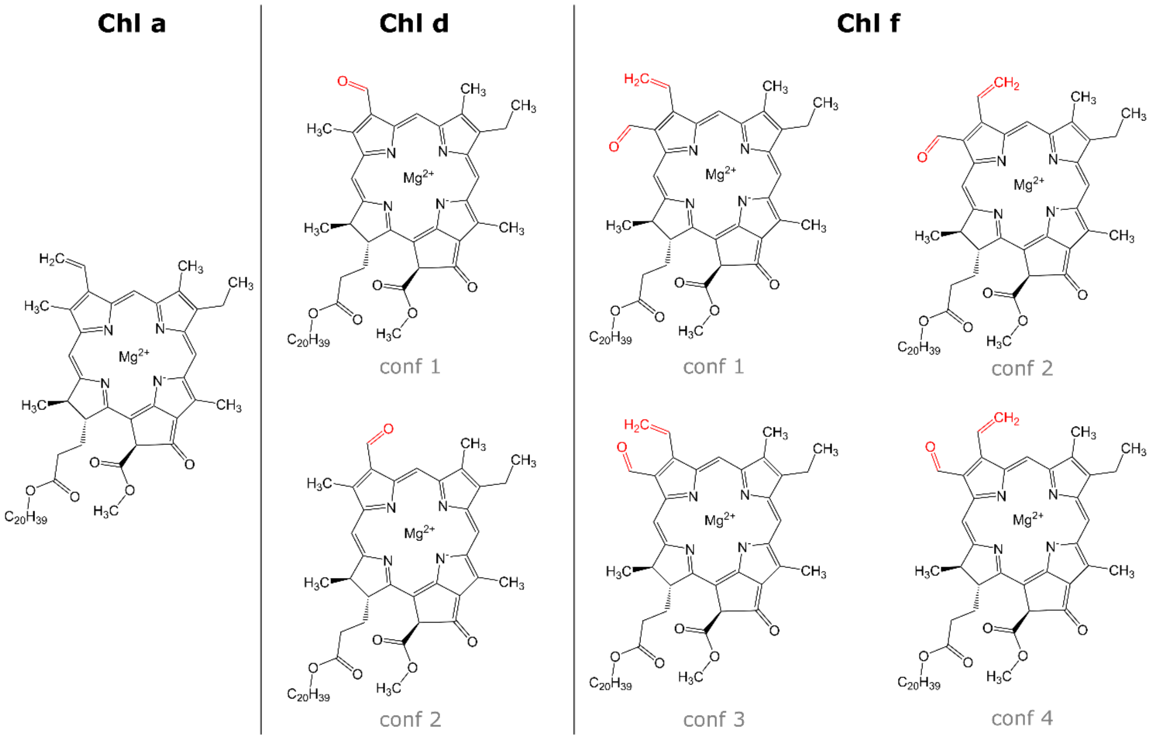


**Figure S9. Structure and different conformations of Chl *a*, Chl *d*, and Chl *f*.** Chl *d* / Chl *f* have two and four rotameric configurations with respect to the formyl group and vinyl groups on C3 and C3, respectively, that affect the hydrogen-bonding in the FRL-PSII.

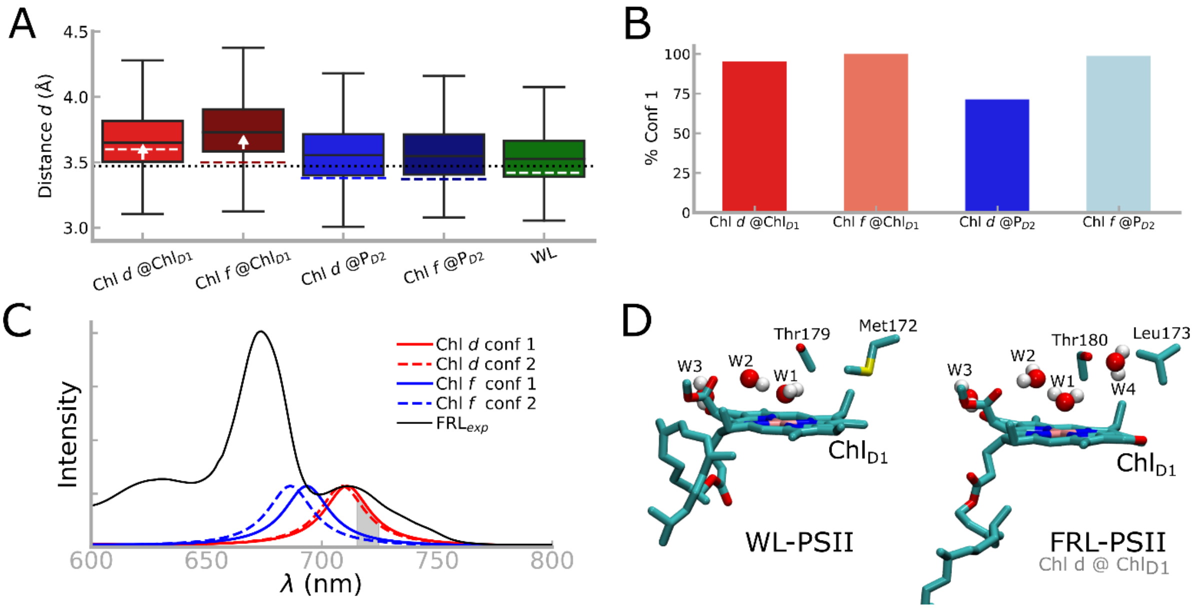


**Figure S10. Conformational dynamics of Chl *d* / *f*.** (A) Inter-pigment spacing between $P_{D1}$ and $P_{D2}$ in our MD simulations. The corresponding QM/MM optimized distances are drawn as vertical dashed lines for each simulation model, while the experimental values for the WL-PSII (3WU2; 3.48 Å) and FRL-PSII (7SA3; 3.46 Å) are shown as a reference as a continuous dotted black line. (B) Distribution of rotamers during the MD simulation. Conf 1 (see Figure S9 for structures) is favored in all simulations. For Chl *d* the oxygen of the formyl group in conf 1 is pointing towards D1-Tyr120/D2-Leu205 (Chl *d* @ Chl$_{D1}$/$P_{D2}$), while for Chl *f*, the oxygen of the formyl group faces away from D1-Tyr120/D2-Leu205 (Chl *d* @ Chl$_{D1}$/$P_{D2}$). Chl *f* lacks a second rotameric conformation in both the Chl *f* @ Chl$_{D1}$ and Chl *f* @ $P_{D2}$ models. The QM/MM optimizations support similar trends, except for Chl *f* @ $P_{D2}$, where conf 3 is favored. (C) Effect of the rotameric state on the spectra (TDDFT level), computed for the isolated Chl$_{D1}$ pigment with D1-Tyr120 and D1-Thr155. The rotameric effect on the $Q_y$ band is small for Chl *d*, but it leads to a spectral shift for Chl *f*. (D) Changed in the water network around the Chl$_{D1}$ site. The smaller sidechain of D1-Leu173 in the FRL-PSII (relative to D1-Met172), allows for additional hydration (W4) in the FRL-PSII.

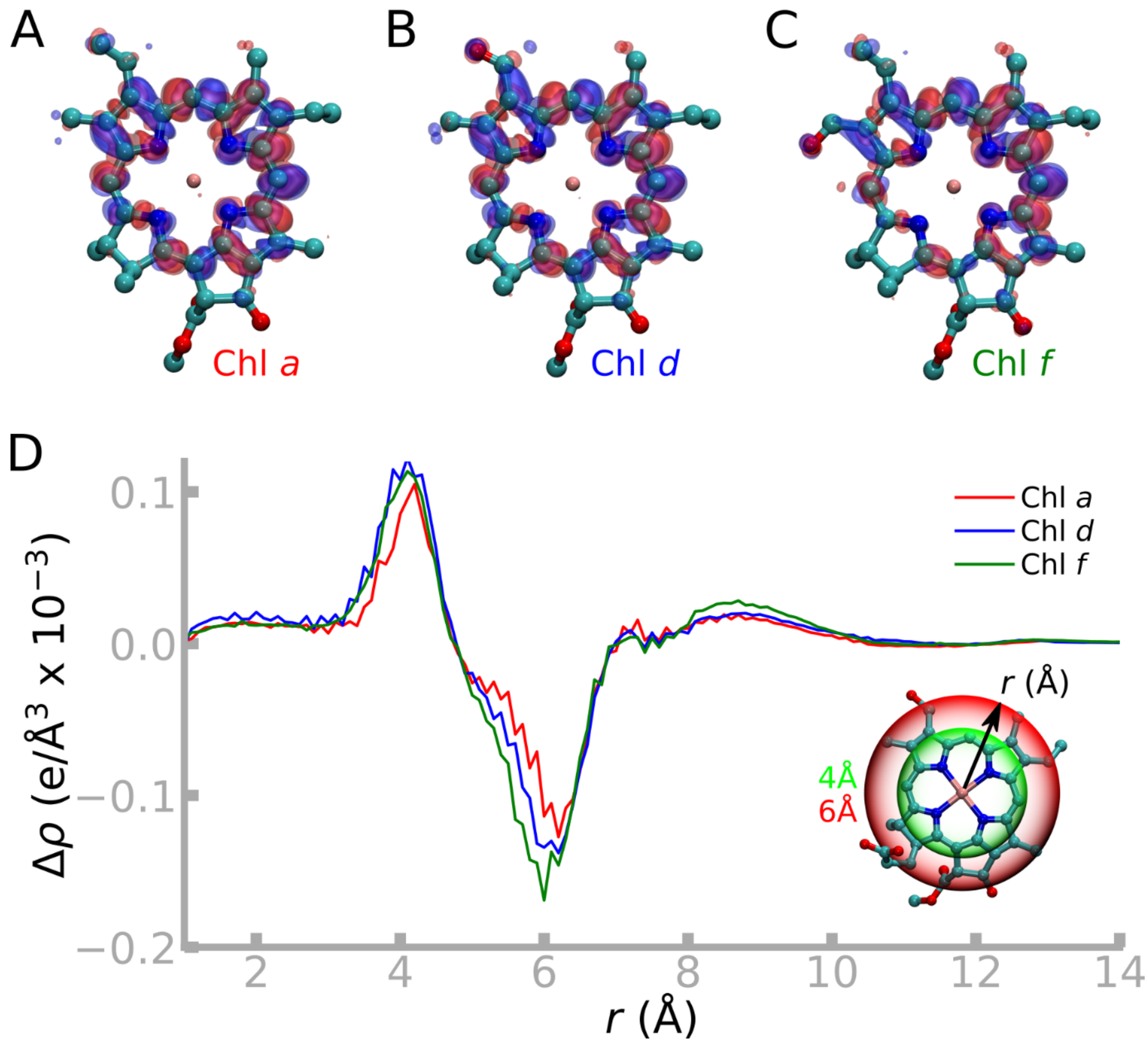


**Figure S11. Density difference upon photo-excitation for different chlorophyll pigments.** Difference density ($S_0$-$S_1$) of the $Q_y$ excited state for (A) Chl *a*, (B) Chl *d*, and (C) Chl *f*. The blue (+0.0005 e/Å$^3$) and red iso-surfaces (-0.0005 e/Å$^3$) represent accumulation and depletion of the electron density upon excitation. (D) Radial excitation energy density difference (Δϱ) as a function of the distance *r* from $Mg^{2+}$ for chlorophyll *a*, *d*, and *f*. The inset shows the radial integration sphere around the chlorophyll center, with surfaces corresponding to the maximum/minimum Δϱ at 4 Å and 6 Å, respectively.

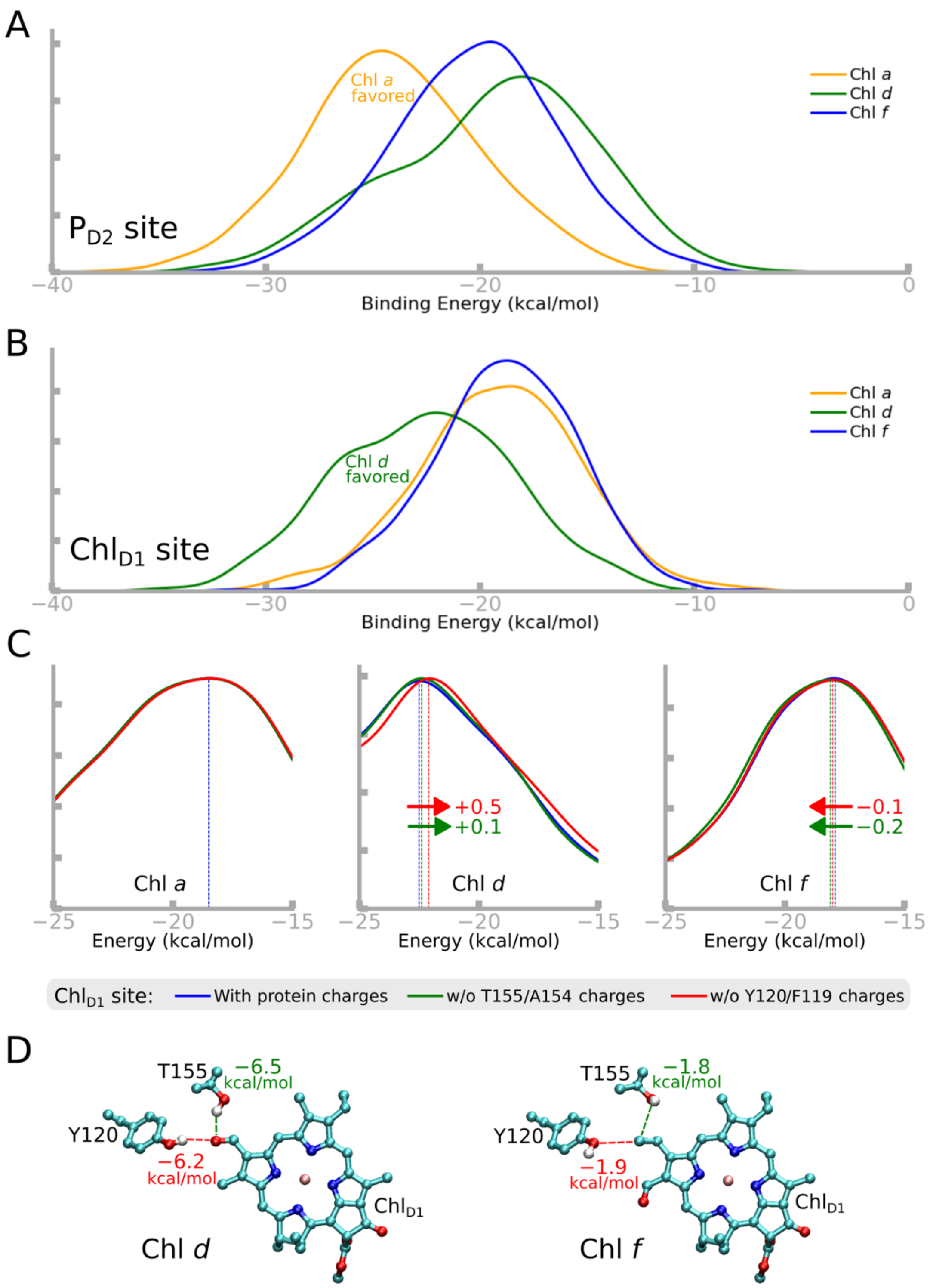


**Figure S12. Binding free energies of chlorophyll clusters in different PSII isoforms** based on MM/PBSA calculations along the MD trajectories (Table S1; see *Extended Methods*). (A) Binding energies for the $Chl_{D1}$ position for Chl *a* in the WL-PSII and Chl *d*/Chl *f* in the FRL-PSII. Chl *d* at the $Chl_{D1}$ position binds the strongest. (B) Binding energies for the $P_{D2}$ position for Chl *a* in the WL-PSII and Chl *d*/Chl *f* in the FRL-PSII system, suggesting that Chl *a* at the $P_{D2}$ position binds the strongest. (C)-(D) Binding energies decomposed into contributions from D1-Thr-155 and D1-Tyr-120 were calculated by (C) switching off the atomic partial charges of the sidechain, or (D) removing the residues from the QM/MM calculations.

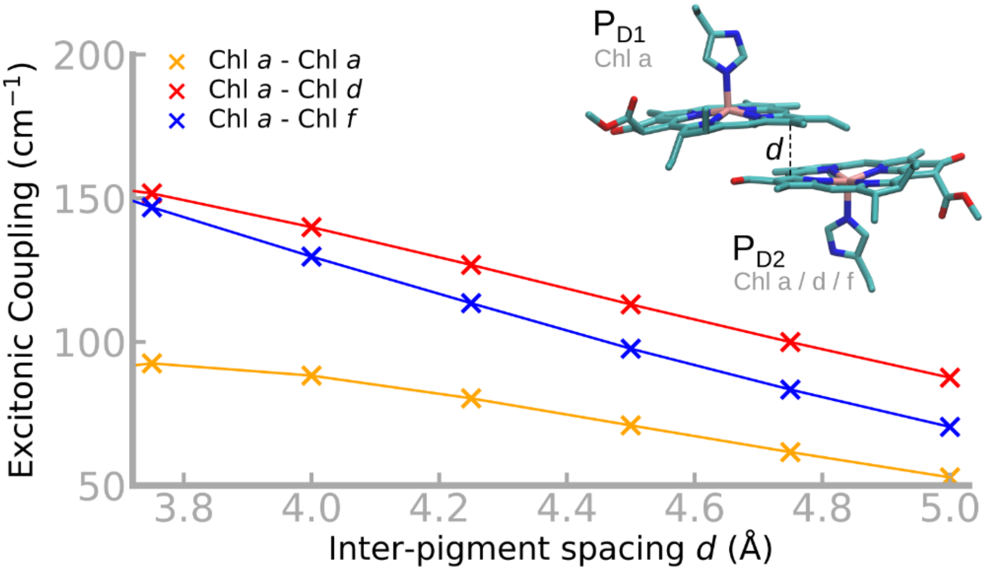


**Figure S13. Excitonic couplings within $P_{D1}$/$P_{D2}$.** Distance dependence of the excitonic couplings within the $P_{D1}$/$P_{D2}$ cluster, with different $P_{D2}$ pigment types. The values shown are obtained at the fragment excitation difference /Tamm-Dancoff approximation (FED/TDA) at the B3LYP/def2-TZVP level, whilst the FED/RPA ansatz yields similar trends but around 30% smaller coupling elements. See Table S11 for further details.

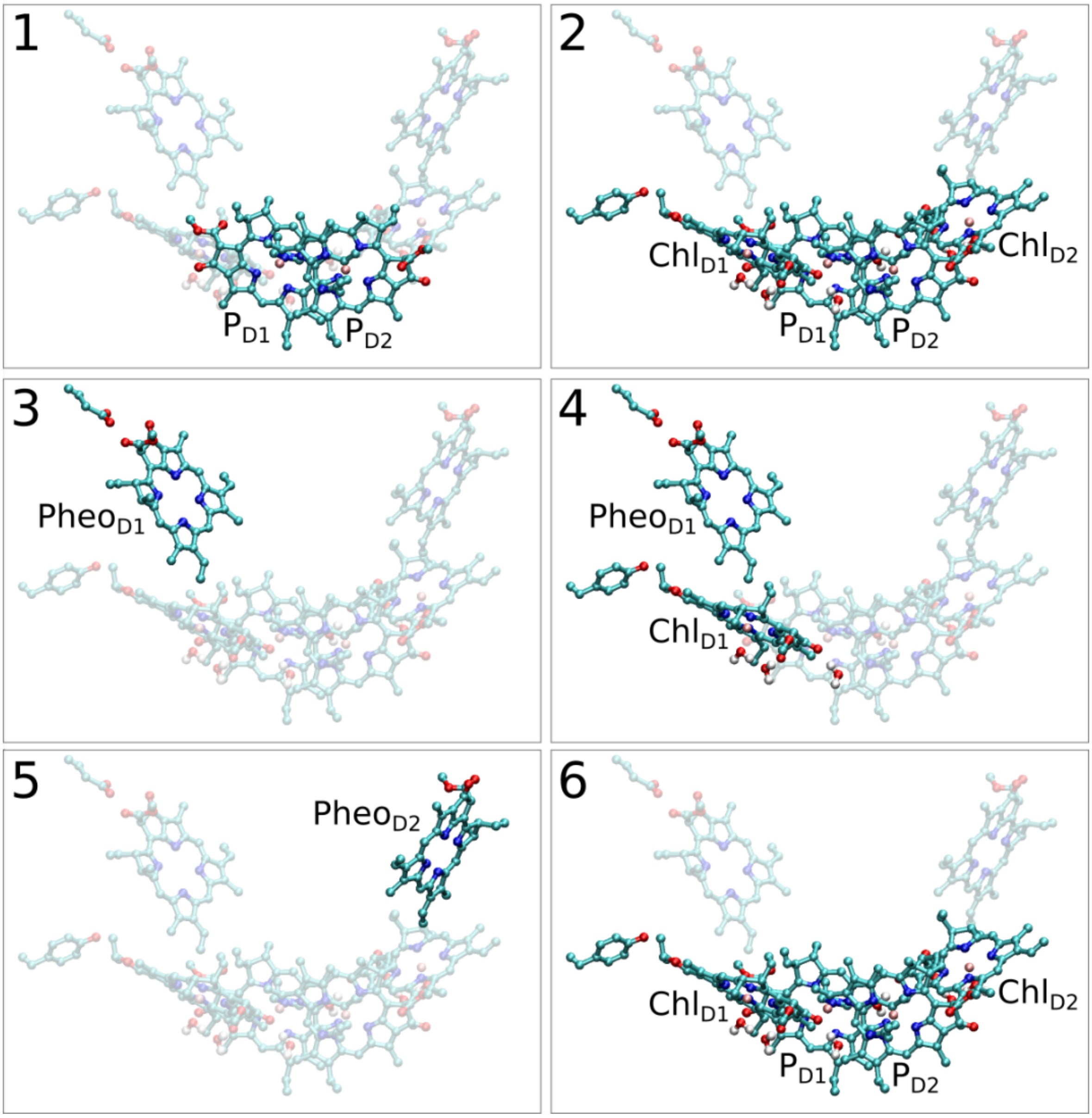


**Figure S14. The micro-iterative optimization scheme of the QM/MM models.** The models were optimized in six sub-steps 1-6 (see *Extended Methods*).

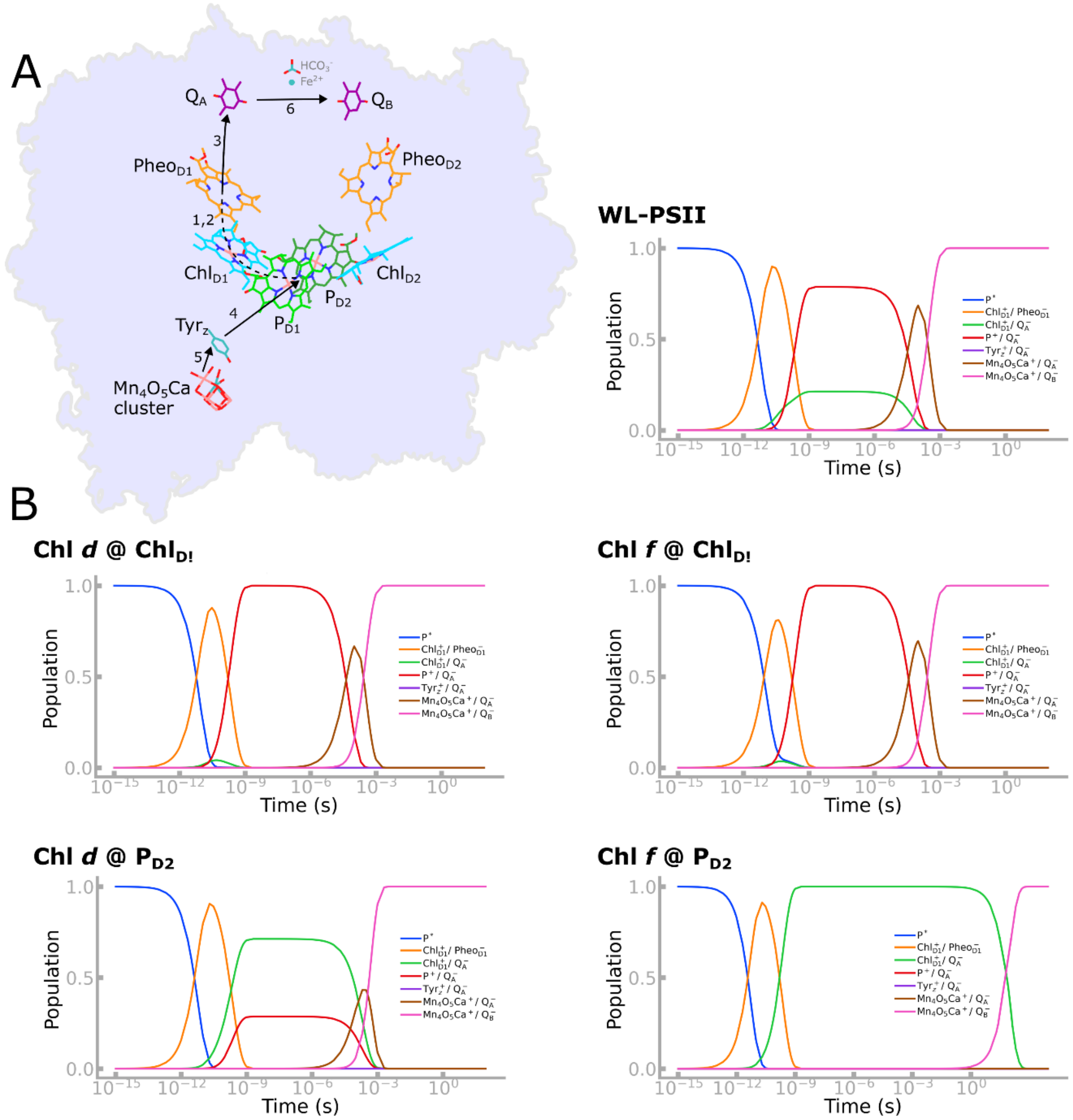


**Figure S15. Comparison of simulated charge transfer kinetics in the WL-PSII and FRL-PSII.** (A) Light-driven charge separation in PSII studied in kinetic models. (B) Charge transfer kinetics for WL-PSII, and FRL-PSII with Chl *d* at $Chl_{D1}$ (Chl *d* @ $Chl_{D1}$); FRL-PSII with Chl *f* at $Chl_{D1}$ (Chl *f* @ $Chl_{D1}$); FRL-PSII with Chl *d* at $P_{D2}$ (Chl *d* @ $P_{D2}$); and FRL-PSII with Chl *f* at $P_{D2}$ (Chl *f* @ $P_{D2}$).

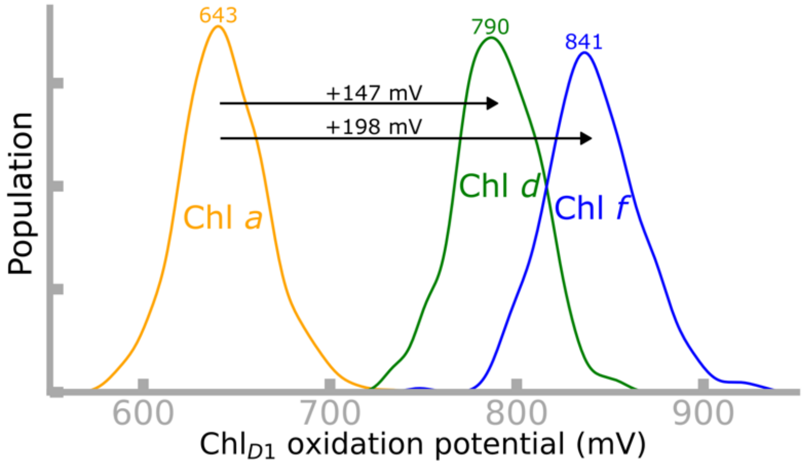


**Figure S16**. **Calculated oxidation potentials.** Oxidation potential for $Chl_{D1}$ (in the presence of $Pheo_{D1}^{+}$) with Chl *a* (WL-PSII) or Chl *d*/*f* (FRL-PSII) modeled at the $Chl_{D1}$ site. The calculations are based on MD simulations of the respective constructs followed by PBC/MC calculations (see *Extended Methods*).

**Table S1. List of QM/MM models and MD simulations.** For the FRL-PSII models, the $P_{D2}$ and $Chl_{D1}$ positions were exchanged with Chl *a*/*d*/*f* pigments, whereas $P_{D1}$ and $Chl_{D2}$ were exclusively modeled with Chl *a* in all models. The C508 pigment pocket in CP43 was exchanged with Chl *d* or Chl *f* based on Ref. 39.

| **Model nomenclature** | **System** | **$P_{D2}$** | **$Chl_{D1}$** | **C508** | **QM/MM optimization** | **Simulation time (MD)** |
|---|---|---|---|---|---|---|
| Chl *d* @ $Chl_{D1}$ | FRL | Chl *a* | Chl *d* | Chl *f* | 6 substeps each (see "*Extended Methods* - QM/MM models") | 2 x 300 ns |
| Chl *f* @ $Chl_{D1}$ | | Chl *a* | Chl *f* | Chl *d* | | 2 x 300 ns |
| Chl *d* @ $P_{D2}$ | | Chl *d* | Chl *a* | Chl *f* | | 2 x 300 ns |
| Chl *f* @ $P_{D2}$ | | Chl *f* | Chl *a* | Chl *d* | | 2 x 300 ns |
| WL | WL | Chl *a* | Chl *a* | Chl *a* | | 2 x 300 ns |
| | | | | | | **Total: 3 μs** |

**Table S2. $D_1$ diagnostics for RVS-LT-SOS-ADC(2) calculations of different chlorophylls types.** The calculations were performed in the gas-phase and in the protein environment.

| **Pigment at $Chl_{D1}$ position** | **Model** | **$D_1$** | |
|---|---|---|---|
| | | Gas phase | Protein environment |
| Chl *a* | WL | 0.061 | 0.061 |
| Chl *d* | Chl *d* @ $Chl_{D1}$ | 0.059 | 0.056 |
| Chl *f* | Chl *f* @ $Chl_{D1}$ | 0.057 | 0.057 |

**Table S3. Benchmarking VEEs for Chl *a*.** The calculations were performed at the RVS-LT-SOS-ADC(2) and TDDFT levels and compared against experimental gas-phase data.[40] The first five excited states ($S_1$-$S_5$), spanning across the Q- and B-bands, along with VEEs (in eVs) and oscillator strengths ($f_{osc}$) are shown. The canonical orbitals involved in the excitations are also marked (H - HOMO; L - LUMO). The experimental values shown in the table refer to band-maxima.

| State Assignment | | 1 | 2 | - | 3 | - | 4 |
|---|---|---|---|---|---|---|---|
| | | $Q_y$ | $Q_x$ | | $B_x$ | | $B_y$ |
| **Experimental** | *E* (eV)* | 1.94 | 2.23 | - | 3.08 | - | 3.38 |
| **RVS-LT-SOS-ADC(2)** | *E* (eV) | 1.96 | 2.35 | - | 3.21 | - | 3.43 |
| | $f_{osc}$ | 0.25 | 0.03 | | 1.08 | | 0.93 |
| **ωB97X-V** | *E* (eV) | 1.92 | 2.49 | - | 3.44 | 3.64 | 3.80 |
| | $f_{osc}$ | 0.23 | 0.03 | | 1.01 | 0.35 | 0.95 |
| **ωB97M-V** | *E* (eV) | 1.92 | 2.45 | - | 3.40 | 3.60 | 3.75 |
| | $f_{osc}$ | 0.22 | 0.03 | | 0.98 | 0.34 | 0.94 |
| **LRC-ωPBEh** | *E* (eV) | 2.01 | 2.39 | - | 3.34 | 3.45 | 3.63 |
| | $f_{osc}$ | 0.23 | 0.03 | | 0.73 | 0.37 | 0.93 |
| **CAM-B3LYP** | *E* (eV) | 2.03 | 2.40 | - | 3.34 | 3.42 | 3.62 |
| | $f_{osc}$ | 0.25 | 0.02 | | 0.75 | 0.35 | 0.93 |
| **CAMh-B3LYP** | *E* (eV) | 2.06 | 2.36 | 3.25 | 3.30 | - | 3.50 |
| | $f_{osc}$ | 0.25 | 0.02 | 0.22 | 0.72 | | 0.69 |

*Band maximum.

**Table S4. Calculated excitation energies of the FRL-PSII and WL-PSII.** The table shows vertical excitation energies at the ωB97X-V and RVS-LT-SOS-ADC(2) levels (unshifted). The corresponding spectra are shown in main text Figure 2.

| | | | **Experimental** | | **ωB97X-V** | | | **RVS-LT-SOS-ADC(2)** | | |
|---|---|---|---|---|---|---|---|---|---|---|
| | | | *E* (eV) | λ (nm) | *E* (eV) | λ (nm) | $f_{osc}$ | *E* (eV) | λ (nm) | $f_{osc}$ |
| FRL | 674 nm peak | Chl *d* @ $Chl_{D1}$ | 1.840 | 674 | 1.902 | 652 | 0.045 | 2.025 | 640 | 0.012 |
| | | Chl *f* @ $Chl_{D1}$ | | | 1.898 | 653 | 0.060 | 2.037 | 635 | 0.011 |
| | | Chl *d* @ $P_{D2}$ | | | 1.952 | 635 | 0.038 | 2.068 | 626 | 0.012 |
| | | Chl *f* @ $P_{D2}$ | | | 1.949 | 636 | 0.035 | 2.064 | 627 | 0.012 |
| | far-red peak | Chl *d* @ $Chl_{D1}$ | 1.734 to 1.710 | 715 to 725 | 1.812 | 684 | 0.038 | 1.967 | 661 | 0.012 |
| | | Chl *f* @ $Chl_{D1}$ | | | -* | - | - | 1.955 | 664 | 0.011 |
| | | Chl *d* @ $P_{D2}$ | | | 1.782 | 696 | 0.028 | 1.962 | 663 | 0.007 |
| | | Chl *f* @ $P_{D2}$ | | | 1.793 | 691 | 0.039 | 1.872 | 694 | 0.007 |
| WL | 674 nm peak | | 1.840 | 674 | 1.933 | 641 | 0.059 | 2.056 | 631 | 0.015 |

* For Chl *f* @ $Chl_{D1}$ the 674 nm and the far-red peak merge for ωB97X-V, therefore no values are reported here.

**Table S5. Summary of VEEs for the FRL-PSII models from ADC(2) calculations.** The table summarizes site energies ($Q_y$ excitation) and corresponding oscillator strengths ($f_{osc}$) of individual pigments for the FRL-PSII models. The excited state computations were performed at the RVS-LT-SOS-ADC(2)/def2-TZVP level of theory. The lowest $Q_y$ states of each model are highlighted. Corresponding TDDFT values are listed in Table S6.

| | | **Chl *d* @ Chl$_{D1}$** | **Chl *f* @ Chl$_{D1}$** | **Chl *d* @ P$_{D2}$** | **Chl *f* @ P$_{D2}$** | **WL** |
|---|---|---|---|---|---|---|
| P$_{D1}$/P$_{D2}$ | *E* (eV) | 1.936 | 1.943 | 1.867 | 1.786 | 1.946 |
| | $f_{osc}$ | 0.335 | 0.348 | 0.312 | 0.361 | 0.285 |
| P$_{D1}$ | *E* (eV) | 1.977 | 1.983 | 1.979 | 1.978 | 1.982 |
| | $f_{osc}$ | 0.253 | 0.252 | 0.239 | 0.235 | 0.232 |
| P$_{D2}$ | *E* (eV) | 2.002 | 2.006 | 1.919 | 1.836 | 2.023 |
| | $f_{osc}$ | 0.245 | 0.247 | 0.277 | 0.363 | 0.223 |
| Chl$_{D1}$ | *E* (eV) | 1.904 | 1.897 | 2.007 | 2.006 | 1.993 |
| | $f_{osc}$ | 0.342 | 0.343 | 0.276 | 0.277 | 0.295 |
| Chl$_{D2}$ | *E* (eV) | 1.978 | 1.987 | 2.007 | 2.011 | 2.009 |
| | $f_{osc}$ | 0.287 | 0.280 | 0.270 | 0.265 | 0.263 |
| Pheo$_{D1}$ | *E* (eV) | 1.956 | 1.992 | 2.014 | 2.012 | 2.064 |
| | $f_{osc}$ | 0.133 | 0.133 | 0.137 | 0.139 | 0.139 |
| Pheo$_{D2}$ | *E* (eV) | 2.133 | 2.138 | 2.073 | 2.084 | 1.994 |
| | $f_{osc}$ | 0.136 | 0.134 | 0.134 | 0.136 | 0.171 |

**Table S6. Summary of VEEs for the FRL-PSII models from TDDFT calculations.** The low-energy $Q_y$ excitations and corresponding oscillator strengths ($f_{osc}$) of individual pigments for the FRL-PSII models. The excited state computations were performed at the LR-TDDFT (ωB97X-V)/def2-TZVP level of theory. The lowest $Q_y$ states of each model are highlighted. Corresponding ADC(2) values are listed in Table S5.

| | | **Chl *d* @ Chl$_{D1}$** | **Chl *f* @ Chl$_{D1}$** | **Chl *d* @ P$_{D2}$** | **Chl *f* @ P$_{D2}$** | **WL** |
|---|---|---|---|---|---|---|
| P$_{D1}$/P$_{D2}$ | *E* (eV) | 1.912 | 1.912 | 1.791 | 1.802 | 1.915 |
| | $f_{osc}$ | 0.325 | 0.335 | 0.236 | 0.327 | 0.286 |
| P$_{D1}$ | *E* (eV) | 1.947 | 1.948 | 1.953 | 1.944 | 1.947 |
| | $f_{osc}$ | 0.214 | 0.214 | 0.210 | 0.211 | 0.200 |
| P$_{D2}$ | *E* (eV) | 1.958 | 1.960 | 1.831 | 1.839 | 1.969 |
| | $f_{osc}$ | 0.208 | 0.209 | 0.230 | 0.308 | 0.200 |
| Chl$_{D1}$ | *E* (eV) | 1.834 | 1.896 | 1.984 | 1.979 | 1.963 |
| | $f_{osc}$ | 0.265 | 0.289 | 0.239 | 0.238 | 0.251 |
| Chl$_{D2}$ | *E* (eV) | 1.962 | 1.968 | 1.991 | 1.991 | 1.990 |
| | $f_{osc}$ | 0.238 | 0.233 | 0.228 | 0.225 | 0.230 |
| Pheo$_{D1}$ | *E* (eV) | 1.913 | 1.941 | 1.969 | 1.964 | 2.015 |
| | $f_{osc}$ | 0.122 | 0.120 | 0.124 | 0.126 | 0.126 |
| Pheo$_{D2}$ | *E* (eV) | 2.097 | 2.096 | 2.033 | 2.037 | 1.947 |
| | $f_{osc}$ | 0.119 | 0.118 | 0.126 | 0.124 | 0.152 |

**Table S7. VEEs for different pigments at the $Chl_{D1}$ position.** The VEEs ($Q_y$ states, in eV) and corresponding oscillator strength of different pigments at the $Chl_{D1}$ position for all models (Table S1) calculated at the RVS-LT-SOS-ADC(2)/def2-TZVP and ωB97X-V/def2-TZVP levels. The computation was performed for monomeric pigment models ($Chl_{D1}$ + two water molecules), and by probing the effect of specific protein residues. The effect of D1-Phe119/D1-Ala154 (WL-PSII) and D1-Tyr120/D1-Thr155 (FRL-PSII) was studied by switching off the point charges in the QM models. "WL Mut" refers to the WL-PSII model, with F119Y and A154T mutations.

| Pigment type at $Chl_{D1}$ position (model) | Environment | RVS-LT-SOS-ADC(2) | | | ωB97X-V | | |
|---|---|---|---|---|---|---|---|
| | | *E* (eV) | *λ* (nm) | $f_{osc}$ | *E* (eV) | *λ* (nm) | $f_{osc}$ |
| Chl *a* (WL) | Gas phase | 1.971 | 629 | 0.260 | 1.953 | 635 | 0.228 |
| | with protein charges | 1.993 | 622 | 0.295 | 1.963 | 631 | 0.251 |
| | w/o Phe119 charges | 1.994 | 622 | 0.294 | 1.964 | 631 | 0.251 |
| | w/o Ala154 charges | 1.998 | 620 | 0.288 | 1.970 | 629 | 0.248 |
| Chl *a* (WL Mut) | Gas phase | 1.968 | 630 | 0.261 | 1.955 | 634 | 0.226 |
| | With protein charges | 2.008 | 617 | 0.314 | 1.981 | 626 | 0.252 |
| | w/o Tyr119 charges | 2.014 | 616 | 0.313 | 1.987 | 624 | 0.252 |
| | w/o Thr154 charges | 1.994 | 622 | 0.318 | 1.968 | 630 | 0.255 |
| Chl *a* (Chl *d* @ $P_{D2}$) | Gas phase | 1.983 | 625 | 0.250 | 1.971 | 629 | 0.221 |
| | with protein charges | 2.007 | 618 | 0.276 | 1.984 | 625 | 0.239 |
| | w/o Tyr120 charges | 2.015 | 615 | 0.273 | 1.992 | 622 | 0.238 |
| | w/o Thr155 charges | 2.002 | 619 | 0.278 | 1.980 | 626 | 0.240 |
| Chl *d* (Chl *d* @ $Chl_{D1}$) | Gas phase | 1.907 | 650 | 0.283 | 1.843 | 673 | 0.233 |
| | with protein charges | 1.904 | 651 | 0.342 | 1.834 | 676 | 0.265 |
| | w/o Tyr120 charges | 1.921 | 646 | 0.337 | 1.850 | 670 | 0.261 |
| | w/o Thr155 charges | 1.920 | 646 | 0.331 | 1.852 | 670 | 0.257 |
| Chl *f* (Chl *f* @ $Chl_{D1}$) | Gas phase | 1.841 | 674 | 0.296 | 1.841 | 673 | 0.266 |
| | with protein charges | 1.897 | 654 | 0.343 | 1.896 | 654 | 0.289 |
| | w/o Tyr120 charges | 1.885 | 658 | 0.346 | 1.884 | 658 | 0.291 |
| | w/o Thr155 charges | 1.884 | 658 | 0.343 | 1.885 | 658 | 0.289 |

**Table S8. Distribution of charge/spin upon oxidation of $P_{D1}$/$P_{D2}$.** The calculations were performed at the B3-LYP/def2-TZVP level (gas phase).

| **Chl type ($P_{D1}$ + $P_{D2}$)** | **$P_{D1}$** | | **$P_{D2}$** | |
|---|---|---|---|---|
| | Charge (%) | Unpaired $e^-$ (%) | Charge (%) | Unpaired $e^-$ (%) |
| *a* + *a* | 58.8 | 56.6 | 41.2 | 43.4 |
| *a* + *d* | 79.6 | 80.5 | 20.4 | 19.5 |
| *a* + *f* | 80 | 82.1 | 20 | 17.9 |

**Table S9**. **Calculated and experimental reduction and oxidation potentials used as model compounds for Chl *a*, Chl *d*, and Chl *f*.** The experimental values correspond to the isolated chromophore in acetonitrile (AN) [41]. Calculated redox/oxidation potentials were obtained from DFT calculations at B3LYP-D3/def2-TZVP level, and accounting for contributions from zero-point energies, enthalpic, entropic, and solvation energy effects. See *Extended Methods* section for a more detailed description. Only the lowest energy rotameric states of the Chl *d*/*f* formyl group were considered (see Figure S9).

| Chlorophyll type | Ligand | Calc oxidation potential in water (mV) | Calc reduction potential in water (mV) | Calc oxidation potential in AN (mV) | Calc reduction potential in AN (mV) | Exp oxidation potential in AN (mV) | Exp reduction potential in AN (mV) |
|---|---|---|---|---|---|---|---|
| **Chl *a*** | no ligand | 792 | -1166 | 744 | -1235 | 881 | -1120 |
| | His | 410 | -1300 | 368 | -1447 | | |
| | $H_2O$ | 313 | -1203 | 287 | -1330 | | |
| **Chl *d*** (conf 2) | no ligand | 690 | -990 | 663 | -1090 | 880 | -910 |
| | His | 395 | -1015 | 368 | -1154 | | |
| | $H_2O$ | 490 | -1059 | 458 | -1191 | | |
| **Chl *f*** (conf 2) | no ligand | 528 | -933 | 498 | -1008 | 920 | -750 |
| | His | 667 | -1148 | 638 | -1275 | | |
| | $H_2O$ | 584 | -941 | 558 | -1077 | | |

**Table S10. Parameters used for modeling the charge transfer kinetics.** *a* - calculated electrostatic interaction of hole-electron pair, *b* - calculated shifts form PBE/MC models (see Figure S16), *c* - from DFT (B3LYP/def2-TZVP) calculations, *d* - from electron pathway analysis, e - from equation 2 (see *Extended Methods*). $\Delta Chl_{D1}$/ $\Delta P_{D2}$ refers to pigment changes (Chl *d* or *f*) in the $Chl_{D1}$/$P_{D2}$ pigments, with shifts evaluated from PBE/MC calculations. See *Extended Methods* for technical details. The experimental oxidation and reduction potentials used for the calculation of the kinetic model are found in Table S11. f - based on electron transfer theory; g - based on experiments [42]. The $Q_A \rightarrow Q_B$ rate is predicted too fast based on the Moser-Dutton model, suggesting that significant re-organization couples to the process. The experimental rate for this transition was therefore used in the kinetic model.

| **Process** | | **$\Delta G_{DA}$ (mV)** | **$V^{A-D}$ (mV)[a]** | **$\lambda_{DA\text{-}f}$ (meV)[c]** | **$\lambda_{DA\text{-}b}$ (meV)[c]** | **$\langle r_{ij} \rangle$ (Å)[d], $r_{ij}^{edge\text{-}to\text{-}edge}$(Å), $r_{ij}^{max}$ (Å)** | **$\langle \rho \rangle$[d]** | **$k_f$ ($s^{-1}$)[e]** | **$k_b$ ($s^{-1}$)[e]** |
|---|---|---|---|---|---|---|---|---|---|
| **$Chl_{D1}{}^{*} \rightarrow Chl_{D1}^{+}/Pheo_{D1}^{-}$** | WL | -108 (Chl *a*) | | 245 (*a*) | 223 (*a*) | 12.8, 1.9, 24.4 (*a*) | | 2e+11 (*a*) | 2e+4 (*a*) |
| | $\Delta Chl_{D1}$ | -82/135 (Chl *d,f*) | -127 | 195/179 (*d,f*) | 242/244 (*d,f*) | 12.8, 1.9, 24.9 (d) 12.5, 1.9, 24.9 (f) | 0.72 | 1e+11 (*d,f*) | 3e+8/ 5e+9 (*d,f*) |
| | $\Delta P_{D2}$ | -108 (Chl *d,f*) | | 193 (*d,f*) | 237/232 (*d,f*) | 12.6, 1.9, 24.3 (d) 12.6, 1.9, 25.0 (f) | | 2e+11 (*d,f*) | 3e+4 (*d,f*) |
| **$Pheo_{D1}^{-} \rightarrow Q_A^{-}$ ($Chl_{D1}^{+}$)** | - | -408 | 68 | 450 | 386 | 14.9, 3.7, 25.3 | 0.73 | 5e+9 | 0.7 |
| **$Chl_{D1}^{+} \rightarrow P_{D2}^{+}$ ($Q_A^{-}$)** | WL | -20 (*a*) | | 194 (*a*) | 265 (*a*) | 14.0, 1.9, 28.3 (*a*) | | 3e+10 (*a*) | 8e+9 (*a*) |
| | $\Delta Chl_{D1}$ | -46/-263 (*d,f*) | 8 | 209/214 (*d,f*) | 211/188 (*d,f*) | 13.8, 1.9, 27.3 (*d*) 13.9, 1.9, 28.1 (*f*) | 0.73 | 9e+10/ 1e+11 (*d,f*) | 2e+6/ 8e+4 (*d,f*) |
| | $\Delta P_{D2}$ | 6/223 (*d,f*) | | 209/203 (*d,f*) | 213/203 (*d,f*) | 14.1, 1.8, 28.4 (*d*) 13.9, 1.9, 28.3 (*f*) | | 8e+9/ 2e+4 (*d,f*) | 2e+10/ 1e+11 (*d,f*) |
| **$P_{D2}^{+} \rightarrow Tyr_Z^{+}$ ($Q_A^{-}$)** | WL | -165 (*a*) | 10 | 435 (*a*) | 235 | 22.6, 13.1, 34.6 (*a*) | 0.66 | 2e+4 (*a*) | 2 (*a*) |
| | $\Delta P_{D2}$ | -191/-408 (*d,f*) | | 435/425 (*d,f*) | | 22.2, 14.0, 31.5 (*d*) 21.7, 13.1, 31.3 (*f*) | | 2e+4/ 6e+4 (*d,f*) | 0.2/ 7e-8 (*d,f*) |
| **$Tyr_Z^{+} \rightarrow Mn_4O_5Ca^{+}$ ($Q_A^{-}$)** | - | -190 | 7 | 402 | 241 | 8.8, 2.7, 14.4 | 0.83 | 2e+11 | 3e+6 |
| **$Q_A^{-} \rightarrow Q_B^{-}$ ($Mn_4O_5Ca^{+}$)** | - | -234 | 3 | 609 | 604 | 17.85, 11.05, 24.27 | 0.77 | 3e+6[f] 3e+3[g] | 0.5[f] 6e-4[f,g] |

**Table S11. Experimental oxidation and reduction potentials of redox active pigments and cofactors in the WL-PSII.**

| Center | $E_m$ (mV) | Reference |
|---|---|---|
| $P^+/P^*$ | -660 | [43, 44] |
| $P/P^+$ | +1240 ($P=P_{D2}$)<br>+1260 ($P=Chl_{D1}$) | [45]<br>[43] |
| $Chl_{D1}$ / $Chl_{D1}^-$ | -570 | [43, 46] |
| $Pheo_{D1}$ / $Pheo_{D1}^-$ | -552 | [47] |
| $Q_A$ / $Q_A^{\bullet-}$ | -144 | [44] |
| $Q_B$ / $Q_B^{\bullet-}$ | +90 | [44] |
| $Tyr_Z$ / $Tyr_Z^{\bullet}$ | +1075 | [43] |
| $Mn_4O_5Ca$ ($S_2/S_1$) | +885 | [43] |

**Table S12. Excitonic couplings of central pigments in the WL-PSII and FRL-PSII.** The values were calculated using the fragment excitation difference (FED) approach along with the Tamm-Dancoff approximation (TDA) or random-phase approximation (RPA) at the ωB97X-V/def2-TZVP level.

| Pigment pair | Pigment type | Model | TDA $V_{ab}$ ($cm^{-1}$) | RPA $V_{ab}$ ($cm^{-1}$) |
|---|---|---|---|---|
| **$P_{D1}$ / $P_{D2}$** | Chl *a* / Chl *a* | WL | 77 | 45 |
| | Chl *a* / Chl *a* | Chl *d* @ $Chl_{D1}$ | 97 | 65 |
| | Chl *a* / Chl *a* | Chl *f* @ $Chl_{D1}$ | 105 | 74 |
| | Chl *a* / Chl *d* | Chl *d* @ $P_{D2}$ | 179 | 134 |
| | Chl *a* / Chl *f* | Chl *f* @ $P_{D2}$ | 192 | 137 |
| **$Chl_{D1}$ / $Pheo_{D1}$** | Chl *a* / Pheo *a* | WL | 121 | 110 |
| | Chl *d* / Pheo *a* | Chl *d* @ $Chl_{D1}$ | 149 | 137 |
| | Chl *f* / Pheo *a* | Chl *f* @ $Chl_{D1}$ | 125 | 113 |
| | Chl *a* / Pheo *a* | Chl *d* @ $P_{D2}$ | 106 | 97 |
| | Chl *a* / Pheo *a* | Chl *f* @ $P_{D2}$ | 109 | 99 |

**Table S13. RVS screening for LT-SOS-ADC(2) calculations of Chl *d* at the $Chl_{D1}$ position.** The calculations were performed with the protein environment. All values were compared to the values obtained without RVS.

| | RVS-LT-SOS-ADC(2) | | | |
|---|---|---|---|---|
| **RVS value** | E (eV) | Δ E (%) | $f_{osc}$ | Δ $f_{osc}$ (%) |
| none | 1.905 | 0.0% | 0.321 | 0.0% |
| 100 | 1.924 | 1.0% | 0.322 | 0.0% |
| 90 | 1.934 | 1.6% | 0.320 | 0.0% |
| 80 | 1.948 | 2.3% | 0.316 | 0.3% |
| 70 | 1.950 | 2.4% | 0.319 | 0.1% |
| 60 | 1.954 | 2.6% | 0.325 | 0.2% |
| 50 | 1.980 | 4.0% | 0.330 | 0.5% |
| 45 | 1.988 | 4.4% | 0.336 | 0.8% |
| 40 | 1.994 | 4.7% | 0.342 | 1.1% |
| 35 | 2.026 | 6.4% | 0.349 | 1.5% |
| 30 | 2.133 | 12.0% | 0.349 | 1.5% |
| 20 | 2.133 | 12.0% | 0.397 | 4.0% |
